# Non-Additive Transcriptional Regulation Underpins Additive Survival Trait In Populations Of *Drosophila* Selected For Increased Post Infection Survival

**DOI:** 10.64898/2026.01.27.702193

**Authors:** Tsering Choton, Nidhi Krishna Shrivastava, Harisankar Durga, Aindrila Das, N.G. Prasad

**Affiliations:** Department of Biological Sciences, Indian Institute of Science Education and Research (IISER) Mohali, Sector 81, SAS Nagar, Mohali, Punjab – 140306, India

**Author notes:** Corresponding Author: Nagaraj Guru Prasad.

**Keywords:** Polygenic adaptation, sex-specific regulation, hybrid mis-expression, *Drosophila*, immune response, experimental evolution

## Abstract

Understanding how immunity evolves requires integrating organismal resistance with the underlying genetic and regulatory mechanisms. Using *Drosophila melanogaster* populations experimentally selected for increased survival post infection with *Enterococcus faecalis*, we dissected the quantitative genetic basis of post-infection survival and the transcriptional architecture associated with it. F1 hybrid analyses revealed an additive genetic basis for survivorship in both sexes, consistent with a polygenic response to selection. In contrast, gene expression showed extensive sex-specific regulation and hybrid misexpression. Males strongly upregulated antimicrobial peptides yet suffered greater mortality, suggesting a costly or inefficient immune strategy. Females relied on distinct effector pathways and exhibited superior survivorship. Notably, transcriptional induction of the melanization precursor *PPO1* did not correspond to increased phenoloxidase enzyme activity, revealing a post-transcriptional constraint on immune deployment. Together, these results show that while the genetic basis of survivorship is additive, the evolved immune phenotype is shaped by sex-specific regulatory divergence, hybrid incompatibilities, and functional bottlenecks. This emphasizes the need to integrate quantitative genetic, transcriptomic, and functional assays to understand the evolution of immune defense.

## 1. Introduction

Natural populations are continuously exposed to selection pressures from pathogens, driving genetic and phenotypic diversity in immune defense mechanisms (Lazzaro & Little, 2009a). Although evolutionary theory traditionally posits that immune traits should be under stabilizing selection due to trade-offs with other life-history traits, accumulating empirical evidence suggests that immune traits may often evolve under positive directional selection (Zuk & Stoehr, 2002). The balance between these two forces depends on various ecological and physiological factors, including fluctuating pathogen pressure and the energetic costs associated with immune activation (Zuk & Stoehr, 2002). Trade-offs may arise because investment in immune function diverts energy from other vital processes such as reproduction and growth (Van Der Most et al., 2011; Ardia et al., 2012). However, such trade-offs can be masked by condition-dependent immunity, and in some contexts, non-immunological defenses may buffer against infection without triggering costly immune activation (Parker et al., 2011). The adaptive evolution of immune traits relies fundamentally on the heritability of quantitative traits and the variance in their fitness consequences (Mousseau & Roff, 1987). Insects like *Drosophila melanogaster*, which rely on innate immunity, offer a powerful model to explore these dynamics. The Toll and Imd signalling pathways, in particular, orchestrate the transcriptional regulation of antimicrobial peptides (AMPs), which serve as frontline defenses against Gram-positive and Gram-negative bacteria, respectively (Lemaitre et al., 1997). In addition to AMP production, the phenoloxidase (PO) cascade contributes to host defense through melanization, facilitating wound healing and pathogen encapsulation (Kelly et al., 2018).

Sexual dimorphism further complicates the evolution of immune function (Kelly et al., 2018). Sex differences in immunity are known to vary by immune trait, taxonomic group, study type, and developmental stage (Schmid-Hempel, 2005; Nunn et al., 2009; Klein & Flanagan, 2016). Generally, females exhibit stronger immune responses and greater resistance to infection, a pattern thought to result from life-history trade-offs and divergent reproductive strategies (Zuk & Stoehr, 2002; Schwenke et al., 2016). Males may allocate more resources to mating effort at the cost of immunity, especially in species with intense sexual selection, whereas females who often bear the physical and immunological costs of mating may evolve more robust immune defenses as a counter-strategy (Rolff & Siva-Jothy, 2002; McKean & Nunney, 2008). This sexual asymmetry can drive co-evolutionary dynamics and may be modulated by mating system, pathogen exposure, and hormonal regulation.

In addition, sexually antagonistic selection may promote immune diversity by favoring alleles that are beneficial in one sex but detrimental in the other (Cox & Calsbeek, 2009; Restif & Amos, 2010). Such selection can lead to sex-specific dominance reversals characterised by alleles that are (partially) dominant when expressed in one sex, but (partially) recessive when expressed in the other (Grieshop & Arnqvist, 2018; Grieshop et al., 2024; Geeta Arun et al., 2021). In *Drosophila*, the heterogametic sex (XY) often displays greater phenotypic variability in immune traits, partly due to the sex chromosomes’ contribution to immunity (Kutch & Fedorka, 2015). In general, the Immunocompetence Handicap Hypothesis (ICHH) further suggests that male traits—while costly in terms of immunity—may serve as honest indicators of genetic quality (Folstad & Karter, 1992), offering a potential explanation for persistent sexual dimorphism in immune responses (Nunn et al., 2009).

To dissect the genetic architecture underlying these patterns, experimental evolution offers a valuable framework. Experimental evolution studies in *Drosophila melanogaster* have shown that repeated bacterial infections and selection of survivors across generations lead to evolved resistance and improved post-infection survival (Basu et al., 2024). Such long-term selection appears to act on key immune genes, including *Relish*, *PGRP-LC*, and *Drosomycin*, which regulate pathogen recognition and antimicrobial peptide production (Begun & Whitley, 2000; Maillet et al., 2008). Evidence of adaptive changes in these genes highlights how both recognition and effector components of the immune system evolve under sustained pathogen pressure, reflecting the coevolutionary dynamics between hosts and microbes (Jiggins & Kim, 2005; Hanson et al., 2023). Yet, many questions remain about how these evolved immune traits are inherited, how they interact with sex-specific regulation, and whether they incur physiological costs or constraints. Hybrid genotypes, in particular, offer a powerful means to test hypotheses about dominance, misregulation, and sex-specific expression patterns, especially in immune genes that are under divergent selection in the parental populations. Regulatory incompatibilities in hybrids may reveal underlying epistatic interactions or constraints on immune robustness.

In this study, we have used an experimental evolution approach in *Drosophila melanogaster* to investigate how sex, genetic background, and infection stage influence post-infection survivorship and immune responses following infection with *Enterococcus faecalis*. We combined dominance coefficient estimation, transcriptomic profiling of key immune genes, and enzymatic assays of PO activity across evolved (EE), control (PP), and reciprocal F1 hybrid (EP, PE) genotypes. This integrative framework allows us to examine the extent of sex-specific dominance, the temporal dynamics of AMP and *PPO1* gene expression, and the functional translation of gene expression into immune phenotypes. By doing so, we aim to uncover how immune traits evolve in a sex and genotype-dependent manner and identify potential mismatches between transcriptional and phenotypic outcomes that may reflect regulatory constraints or trade-offs.

## 2. Materials and Methods

### 2.1. Fly regime and maintenance

The populations of *D. melanogaster* used in this study were derived from four Blue Ridge baseline populations (BRB1-4), each a large (adult census size ≈ 2800), outbred population established from the same 19 isofemale lines (Singh et al., 2025). Each replicate of BRB was used to derive three populations that were subjected to three distinct selection regimes for approximately 120 generations (at the time of experiment): E (infected every generation with *Enterococcus faecalis*), P (injury controls), and N (unhandled controls). Thus, the entire experimental evolution study was replicated over four “blocks”, each containing one E, one P, and one N population. Each block (labelled 1-4) was handled on a different day. Each E, P, or N population included ten vials (25 mm diameter × 90 mm height) containing 6-8 ml of standard banana jaggery-yeast food, with eggs collected at a density of approximately 70 eggs per vial. Vials and subsequent cages were reared under standard laboratory conditions (12:12 light: dark cycle, 25°C, 60% relative humidity). The day of egg collection was designated as day 1 of the life cycle, and treatment corresponding to the selection regime was applied on day 12.

For the E populations, on the 12^th^ day post-egg collection, when the flies were 2-3 days old and fully mature, they were sorted into a balanced sex ratio of 200 males and 200 females. The flies were infected by dipping fine pins (0.1 mm, Fine Science Tools, Canada) into a suspension of the gram-positive pathogenic bacterium *E. faecalis* and pricking the dorsolateral surface of the thorax at a specific optical density (OD_600_). After infection, the flies were placed in a plexiglass cage (14 cm length × 16 cm width × 13 cm height) with food in a 90 mm Petri dish. Approximately 40-50% mortality was observed 96 hours post-infection. An oviposition food plate was placed in the cages 18 hours before egg collection, which took place on the 16th day.

The P populations, or the pseudo-infected population, served as the injury control group. In this case, the flies were again sorted into an equal sex ratio of 100 males and 100 females and were pricked with a pin dipped a sterile 10 mM MgSO4 solution to verify the accuracy of the infection and confirm that the mortality observed in the E-population was due to bacterial infection rather than injury from pricking. After 4 days, eggs were collected for the next generation.

The N populations consisted of non-handled flies, also sorted into a balanced sex ratio of 100 males and 100 females, without any infection. All populations were provided with a fresh food plate on day 14 of their life-cycle.

### 2.2. Bacterial culture

*Enterococcus faecalis* (Ef) is a gram-positive entomopathogenic bacterium with an optimum growth temperature of 37°C (Huycke et al., 2001; Lazzaro et al., 2006). The stocks are maintained as 15% glycerol stock at −80°C. The primary culture was set by inoculating a scrap of stock culture in 10ml of LB (Luria-Bertani-Miller, HiMedia) broth. The primary culture was incubated in an incubator shaker (New Brunswick™ Innova ®42/42R Shaker) set at 37°C and 150 RPM overnight. After 14-16 hours, a secondary culture was established by inoculating 100 µl of primary culture in 10 ml of fresh LB and incubated for 4 hours. Once the media turned turbid, it was centrifuged (25°C, 15 mins, 6000 rpm) to get a bacterial pellet. The supernatant was discarded, and the pellet was resuspended in a sterile 10mM MgSO4 solution to obtain the desired OD (OD_600_= 2). This suspension was used to infect the experimental flies.

### 2.3. Standardisation of experimental flies

Experimental eggs were collected from flies cultured under standardised environmental conditions for one generation without any treatments (infection or pseudo-infection) normally used in the maintenance of E and P populations. This methodology effectively eliminates potential non-genetic parental effects (Clare & Luckinbill, 1985), resulting in “standardised flies”. Eggs were obtained from E and P populations at a density of approximately 70 eggs per vial, with ten vials established for each population reared at the same standard laboratory conditions. The 2-3 day old adults were transferred to plexiglass cages measuring (14x16x13 cm) with fresh food. Experimental eggs were collected from these population cages.

### 2.4. Hybrid Cross setup

To set up reciprocal crosses between E and P populations, we first collected eggs from standardised E and P flies at a density of 70 eggs/vial. Once adult flies started eclosing in these vials about 9 to 10 days after egg collection, we collected them within 6 hours of eclosion (to ensure they were virgin) and housed them in 10 flies each in single-sex vials. For each block, we combined these 100 virgin males and 100 females into Plexiglas cages to set up the following four crosses:

1. E♂ × E♀ (EE)
2. E♂ × P♀ (EP)
3. P♂ × E♀ (PE)
4. P♂ × P♀ (PP)

To generate the F1 progeny, we collected eggs from each of the four crosses at a density of 70 eggs/vial. On the 12th day after egg collection, once all flies from these vials had eclosed, three replicate cages for each cross were set up, each containing 50 males and 50 females and were infected with E. faecalis (OD_600_= 2). Additionally, one sham-infected cage with 50 males and 50 females was established for each cross. Mortality was observed over the following four days, and food plates were changed every alternate day after the cages were set up to measure survivorship post infection.

### 2.5. Gene expression

In this study, we aimed to explore the expression dynamics of four essential immune-related genes—*Dual oxidase* (*duox*), *Diptericin* (*dpt*), *Drosomycin* (*drs*), and *Prophenoloxidase1* (*PPO1*)—across three critical time points: Before infection (BI: serving as the control), 48 hours post-infection, and 96 hours post-infection (HPI) with *E. faecalis*. These genes were selected because of their significant roles in the immune defence mechanisms of *Drosophila melanogaster* against pathogenic infections (Dudzic et al., 2015; Lemaitre et al., 1997; Tleiss et al., 2024). *Duox*-dependent reactive oxygen species (ROS) act as a microbicide at the barrier where these cells contact microorganisms, similar to phagocytic ROS. *Duox* generates extracellular H_2_0_2_, which, with lactoperoxidase, converts SCN− to the antibacterial hypothiocyanate (OSCN−) in mucosal fluids (Kim & Lee, 2014). This Duox-lactoperoxidase system provides a strong antimicrobial defense in mammalian epithelial cells (Gattas et al., 2009; Leto & Geiszt, 2006). Duox-dependent ROS generation plays a crucial role in controlling gut-associated bacteria (Bae et al., 2010; Ha et al., 2005; Kim & Lee, 2014). Diptericin is an essential antimicrobial peptide that actively participates in the immune response of insects, especially in *Drosophila*. Its antibacterial activity, particularly against gram-negative bacteria (Mullinax et al., 2025), highlights its importance in protecting against infections. *Drosomycin* plays a critical role in antifungal defence and is also induced when faced with bacterial challenges, showcasing its dual functionality in innate immunity (Zhang & Zhu, 2009). Finally, *PPO1* encodes an enzyme essential for melanin production, a process that is crucial for pathogen defence, wound healing, and cuticle sclerotisation (Cerenius & Söderhäll, 2021; Marieshwari et al., 2023).

For each cross and sex, adult flies aged 2–3 days were sampled. At each time point (Bi, 48 hours after infection (HPI) and 96 HPI), three whole live flies were collected from each of the three replicate experimental cages per cross within one block (B1). Each group of three flies constituted one biological replicate and was immediately transferred to TRIzol reagent (Sigma, T9424) and stored at −80°C until RNA extraction. From each cage, three independent biological replicates (each comprising three flies) were collected for robust statistical analysis. Total RNA was extracted from each homogenized sample using TRI reagent (Sigma, T9424). The purity and concentration of the isolated RNA were assessed using a NanoDrop 2000 spectrophotometer; a 260/280 ratio near 2.0 was considered indicative of high-quality RNA. Complementary DNA (cDNA) was synthesized from 1000 ng of total RNA using a first-strand cDNA synthesis kit (Thermo Fisher Scientific, Cat. no. K1622). Quantitative PCR was performed on an Applied Biosystems QuantStudio 6 Flex Real-Time PCR System (Cat. no. 4485692) using Maxima SYBR Green qPCR MasterMix (Thermo Fisher Scientific, Cat. no. K0221). Each 10 µl reaction contained 5 µl of SYBR Green MasterMix, 1 µl each of forward and reverse primer (1 µM), 2 µl of nuclease-free water, and 1 µl of diluted (1:5) cDNA template. For each of the three biological replicates, the expression of all target genes and the reference gene (Rp49) was measured in three technical replicates.

Gene expression levels were quantified using the comparative Ct (2^(-ΔΔCt)) method (Livak & Schmittgen, 2001). The cycle threshold (Ct) values were obtained from the fluorescence data. The ΔCt was calculated as the difference between the Ct of the target gene and the Ct of the reference gene (Rp49). The average ΔCt value of the pre-infection (Bi) samples served as the calibrator for each gene. The ΔΔCt was derived by subtracting the calibrator’s ΔCt from the ΔCt of each target sample. The final fold change in gene expression was determined using the formula 2^(-ΔΔCt) and log10 of this was used for subsequent visualisation and statistical analysis.

### 2.6. Phenoloxidase activity

Phenoloxidase (PO) plays a key role in the melanisation process by catalysing the oxidation of phenols into quinones, which subsequently polymerise to form melanin (Binggeli et al., 2014). These copper-dependent enzymes are also involved in oxygen transport and cuticle sclerotisation (Marieshwari et al., 2023). The two main forms, PO1 and PO2, are produced as inactive precursors known as prophenoloxidases (PPOs) or zymogens within crystal cells. They are activated through proteolytic cleavage by serine proteases in the Toll-PO cascade, with cSPH35 and cSPH242 facilitating PPO1 activation (Jin et al., 2024). Phenoloxidases exhibit broad-spectrum bactericidal activity and contribute to bacterial melanisation and aggregation (Zhao et al., 2007). In addition to melanin production, the melanisation reaction generates reactive oxygen species (ROS) and other cytotoxic intermediates that may assist in bacterial encapsulation, formation of protective barriers, and regulation of oxidative stress (Ayres & Schneider, 2008). To evaluate PO activity, we used adult *Drosophila* (both males and females) sampled at three different time points (Bi, 48 HPI and 96 HPI) from three biological replicate across two replicate blocks. PO activity was measured using an L-DOPA assay (Shrivastava et al., 2022).

For the assay, three live adults flies from each time point were homogenised in a cold buffer containing 1X PBS and protease inhibitor (Sigma-Aldrich, S8820). The homogenate was centrifuged at 13,000 rpm for 10 minutes at 4°C, and the supernatant was carefully collected into pre-chilled, pre-labelled Eppendorf tubes. Protein concentration was measured for each sample using the Pierce BCA protein estimation kit (Thermo Scientific, 23227).

To determine PO activity, 50 μl of supernatant containing 10 μg of protein was dispensed into a 96-well plate, and 50 μl of 3 mM L-3,4-dihydroxyphenylalanine (L-DOPA) was added to each well. The enzymatic conversion of L-DOPA to dopachrome, producing an orange colour, was used as a measure of activity. Absorbance was recorded at 492nm to quantify PO activity. Each sample was plated in duplicate, with three replicates per cage for each population.

### 2.7. Fecundity

To assess fecundity, fresh food plates were introduced into cages containing infected flies from all the crosses at two post-infection intervals: 0–12 hours post-infection (HPI) and 24–36 HPI. Eggs laid on the plates were counted after each interval. This was performed for all crosses using three replicate cages per cross and was repeated across four independent experimental blocks.

### 2.8. Statistical analysis

All analyses were performed in R version 4.2.0. Survivorship was analyzed using Cox mixed-effects models (coxme package) with the formula:

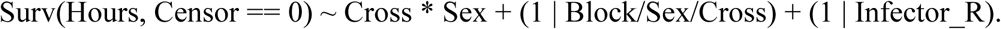

Survivorship was analyzed using Cross and Sex as the fixed factors, with Censor = = 0 indicating death. Block/Sex/Cross interaction, as well as infector identity were modelled as random factors.

We analyzed dominance patterns using a quantitative genetic approach that models survival probability as:

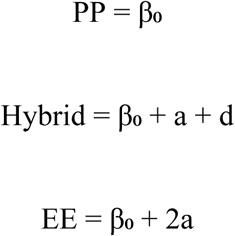

where β₀ represents the baseline survival probability of the PP genotype, “a” is the additive effect per E allele, and “d” is the dominance deviation (Lynch & Walsh 1998). The dominance coefficient “d” was calculated separately for each sex as:

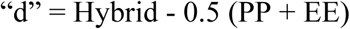

where “Hybrid” represents the mean survival of reciprocal hybrid crosses (EP and PE). Dominance coefficients for survivorship were calculated separately for each sex, with 95% confidence intervals generated via stratified bootstrapping (10,000 iterations).

Gene expression data were analyzed using Kruskal-Wallis tests with Dunn’s post-hoc tests (Benjamini-Hochberg correction). Dominance coefficients for gene expression were calculated for each Gene×Sex×Time combination with confidence intervals determined by bootstrapping (1,000 iterations). Sex differences within Cross×Time combinations used Welch’s t-tests.

PO activation rate (slope of absorbance increase) was analyzed using a linear mixed-effects model with Cross, Sex, and HPI as fixed effects and Block as a random intercept. Type III ANOVA was used to assess the significance of main effects and interactions.

Fecundity was analyzed using a negative binomial generalized linear mixed model (family: nbinom2) with the formula: Eggs_Laid ∼ Cross × Time_Period + (1|Block) + offset(log(Surviving_Females)). This approach accounted for overdispersed count data and varying female numbers across replicates. Significance testing used Wald z-tests, with results reported as estimates ± standard errors and associated p-values.

## 3. Results

### 3.1. Sex-specific dominance in survivorship post-infection with Ef

Post-infection survivorship was significantly influenced by both genetic background and sex (Figure 1; Table 1). The evolved cross (EE) showed the highest survivorship, hybrid crosses (EP and PE) exhibited intermediate survival, and the control cross (PP) showed the lowest survivorship. Across all genetic backgrounds, females consistently survived better than males. Similar survivorship between reciprocal hybrids (EP and PE) indicates that survivorship is not X-linked, but instead reflects an autosomal genetic basis.

**Figure 1.**
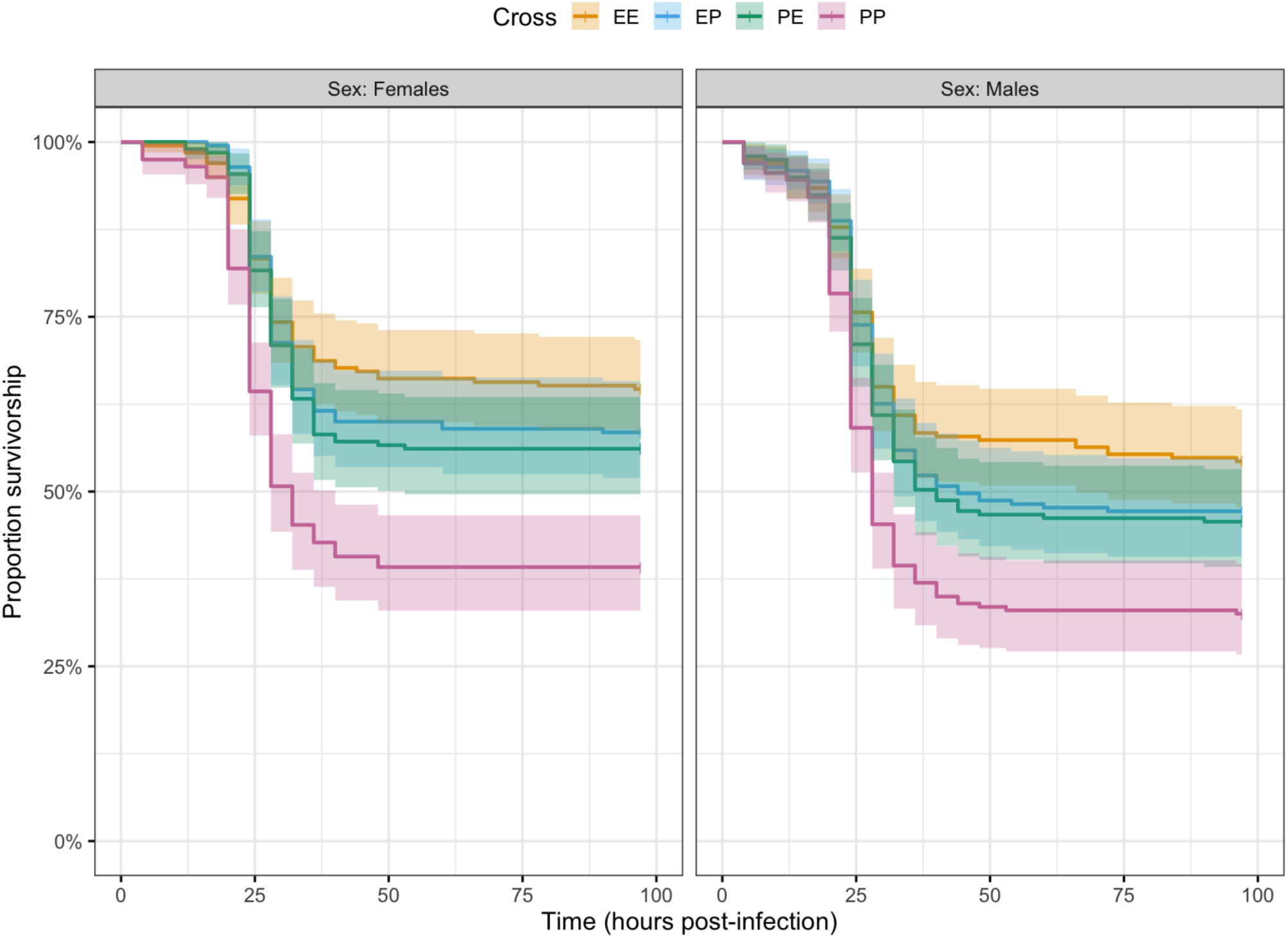
Survival proportions over 96 hours following infection with *E. faecalis* are shown for (a) females and (b) males across EE, EP, PE, and PP F1 crosses. Curves represent block-averaged survival, obtained by fitting survival models within each of the four independent blocks and then aggregating the results across blocks. Block-specific curves are shown in Supplementary Figure S1. A Cox proportional hazards model detected significant effects of Cross and Sex (Table 1).

**Table 1.**
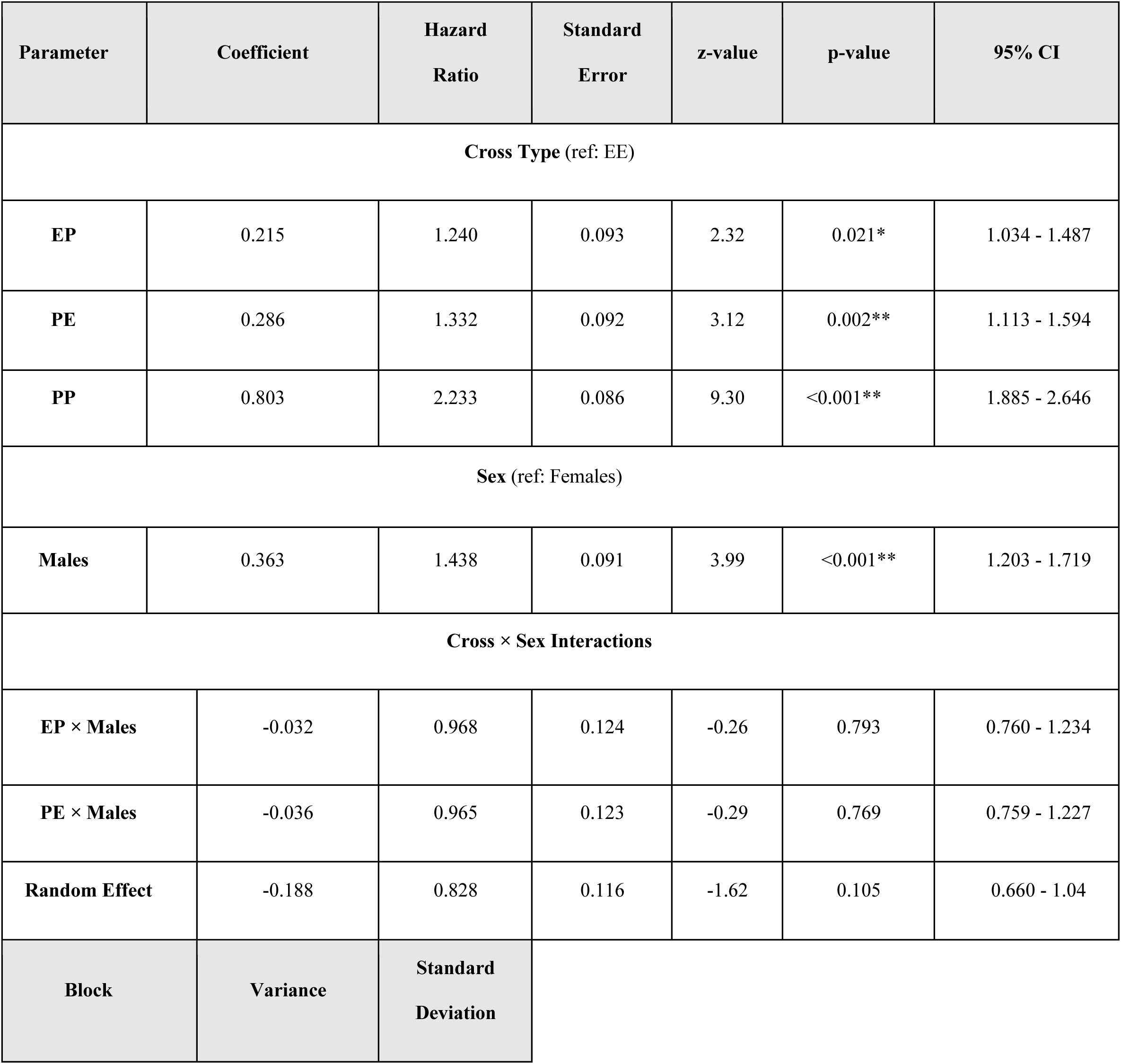
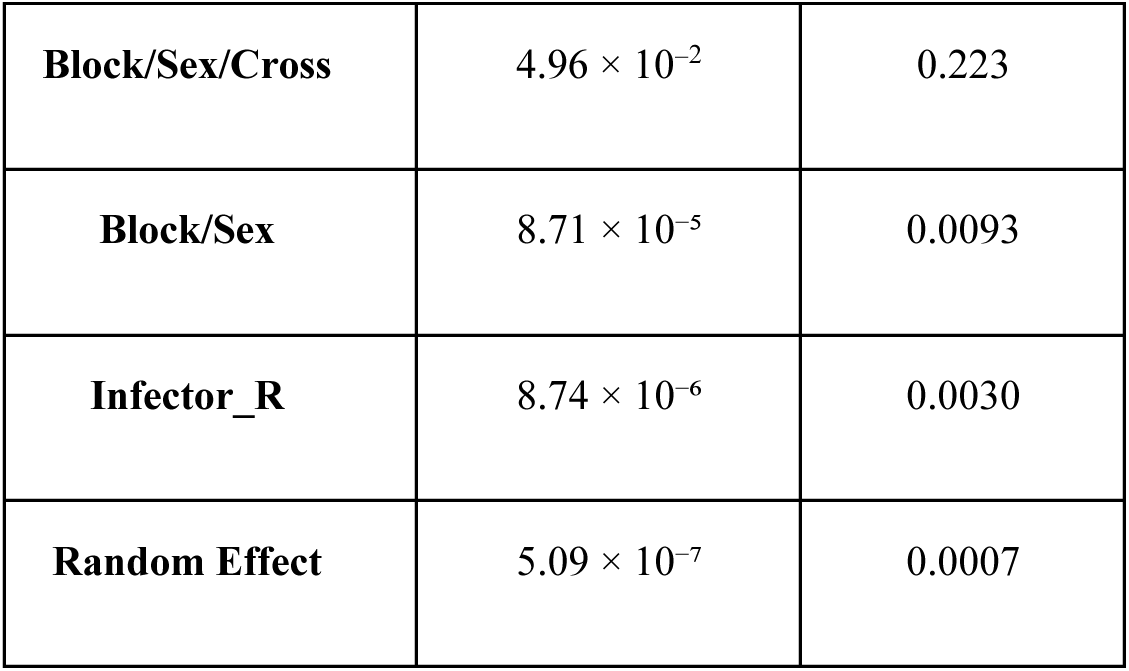
Cox proportional hazards mixed-effects model (coxme) testing effects of cross and sex on post-infection survivorship. The model includes Cross, Sex, and their interaction as fixed effects and Block, Block/Sex/Cross combinations, and Infector as random effects. Reference categories are EE for cross type and Females for sex. Hazard ratios (HR) > 1 indicate increased mortality risk (reduced survivorship), while HR < 1 indicate reduced mortality risk. 95% confidence intervals (CI) are provided. Significant effects: *p < 0.05, **p < 0.01, ***p < 0.001.

Dominance analyses revealed no evidence for sex-specific dominance in post-infection survivorship. Dominance coefficients for females and males were moderate, similar across experimental blocks, and did not differ significantly between sexes (Figure 2; Table 2). Consistent with this, hybrid survivorship closely matched additive expectations based on parental means (Figure 3), indicating that post-infection survivorship is largely additive in its genetic architecture.

**Figure 2.**
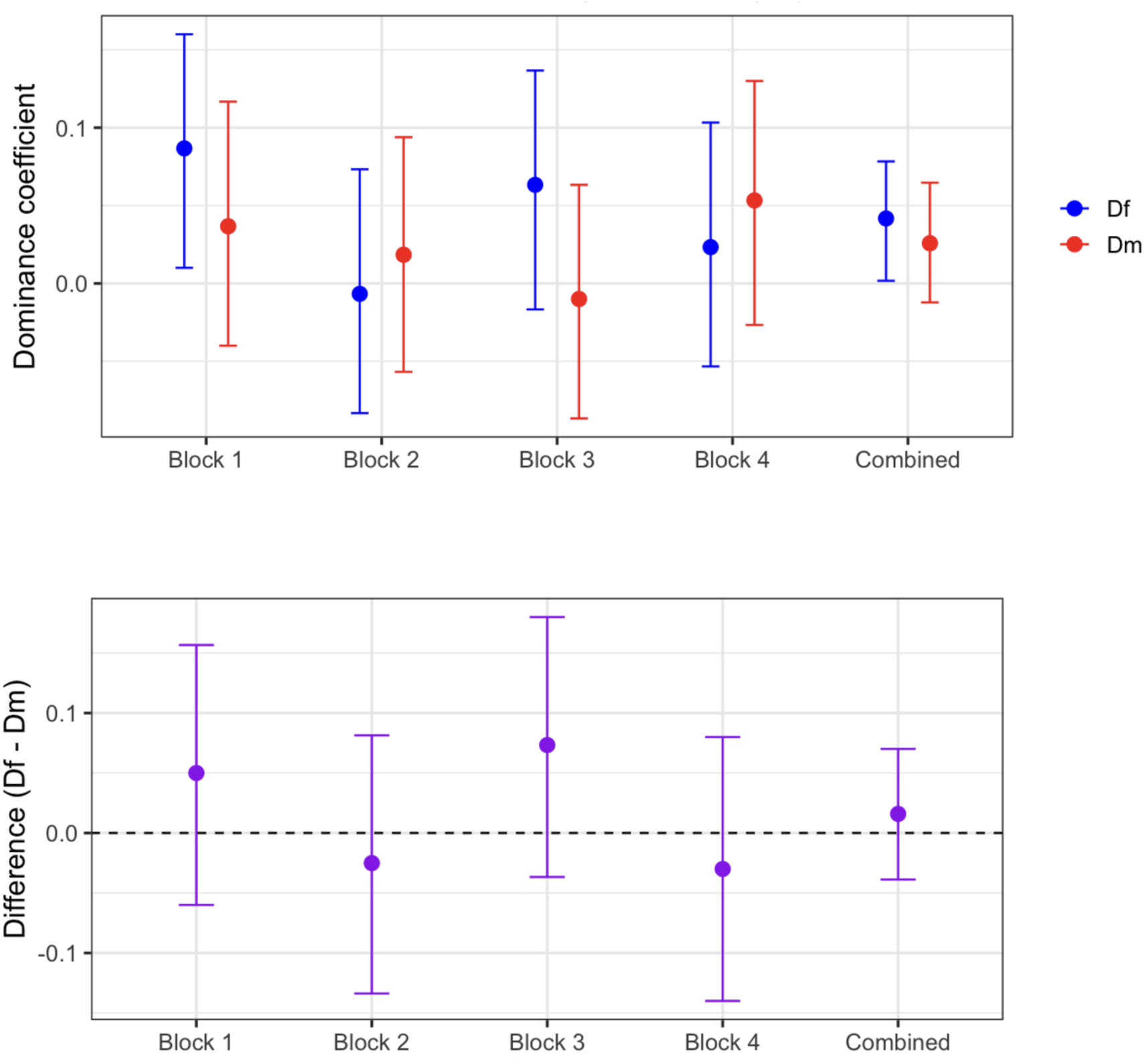
Sex-specific dominance coefficients (Df and Dm) and their differences across experimental blocks. The upper panel shows the estimated dominance coefficients for females (Df, blue) and males (Dm, red) across four replicate blocks and the combined dataset. Error bars represent 95% confidence intervals generated via bootstrap resampling. The lower panel displays the difference between female and male dominance coefficients (Df − Dm) for each block and the combined estimate (purple), with the dashed line at 0 indicating no sex difference. While point estimates suggest moderate dominance in both sexes, the confidence intervals for Df − Dm overlap zero, indicating no consistent or significant sex bias in dominance across blocks.

**Table 2:** Estimated dominance coefficients (Df for females, Dm for males, and their difference Df–Dm) across four replicate blocks and the combined blocks dataset. Values represent point estimates with corresponding lower and upper 95% confidence intervals (CI). A positive Df–Dm indicates higher dominance in females, while a negative value indicates higher dominance in males

| Block | Sex | Value | Lower CI | Upper CI |
| --- | --- | --- | --- | --- |
| Block 1 | Df | 0.0867 | 0.0100 | 0.1600 |
|  | Dm | 0.0367 | -0.0400 | 0.1167 |
|  | Df-Dm | 0.0500 | -0.0600 | 0.1567 |
| Block 2 | Df | -0.0067 | -0.0833 | 0.0733 |
|  | Dm | 0.0184 | -0.0568 | 0.0939 |
|  | Df-Dm | -0.0251 | -0.1338 | 0.0814 |
| Block 3 | Df | 0.0633 | -0.0167 | 0.1367 |
|  | Dm | -0.0100 | -0.0867 | 0.0633 |
|  | Df-Dm | 0.0733 | -0.0367 | 0.1800 |
| Block 4 | Df | 0.0233 | -0.0533 | 0.1033 |
|  | Dm | 0.0533 | -0.0267 | 0.1300 |
|  | Df-Dm | -0.0300 | -0.1400 | 0.0800 |
| Combined | Df | 0.0417 | 0.0017 | 0.0783 |
|  | Dm | 0.0258 | -0.0122 | 0.0647 |
|  | Df-Dm | 0.0159 | -0.0388 | 0.0701 |

**Figure 3.**
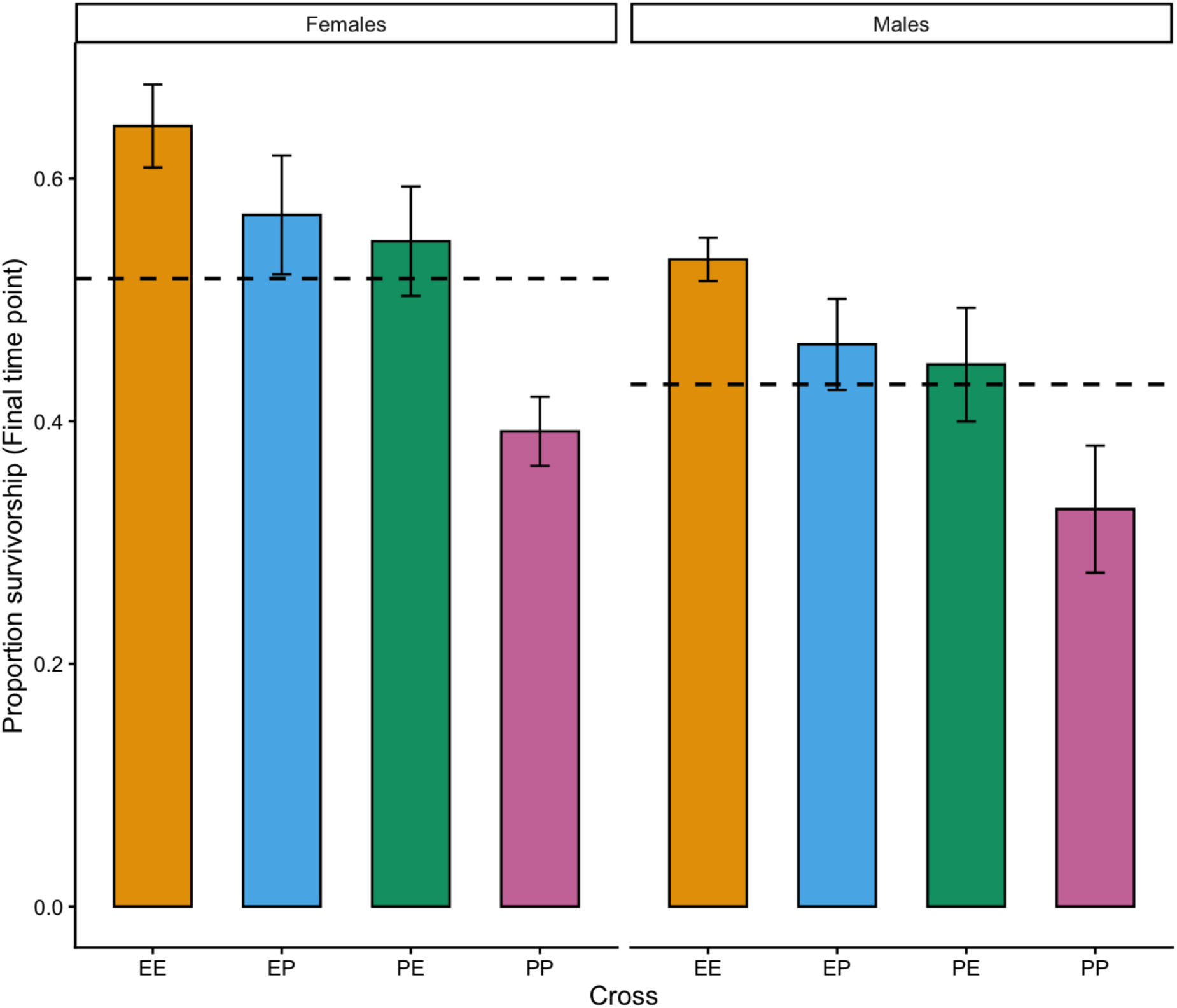
Block-averaged survivorship for each cross (EE, EP, PE, PP) is shown for females and males. Final survivorship was first estimated independently within each block, then averaged across blocks. Dashed lines denote sex-specific additive expectations (mid-parent means). Both hybrid crosses exceed their additive expectations, indicating partial dominance of alleles influencing post-infection survivorship.

### 3.2. Gene Expression

#### 3.2.1 Dual Oxidase (duox) exhibits female-biased expression with late-stage male-specific genetic effects

*Duox* expression consistently displayed a strong female-biased pattern across genetic backgrounds and time points, with large effect sizes indicating biologically meaningful sex differences (Figure 4, Table 3).

**Figure 4.**
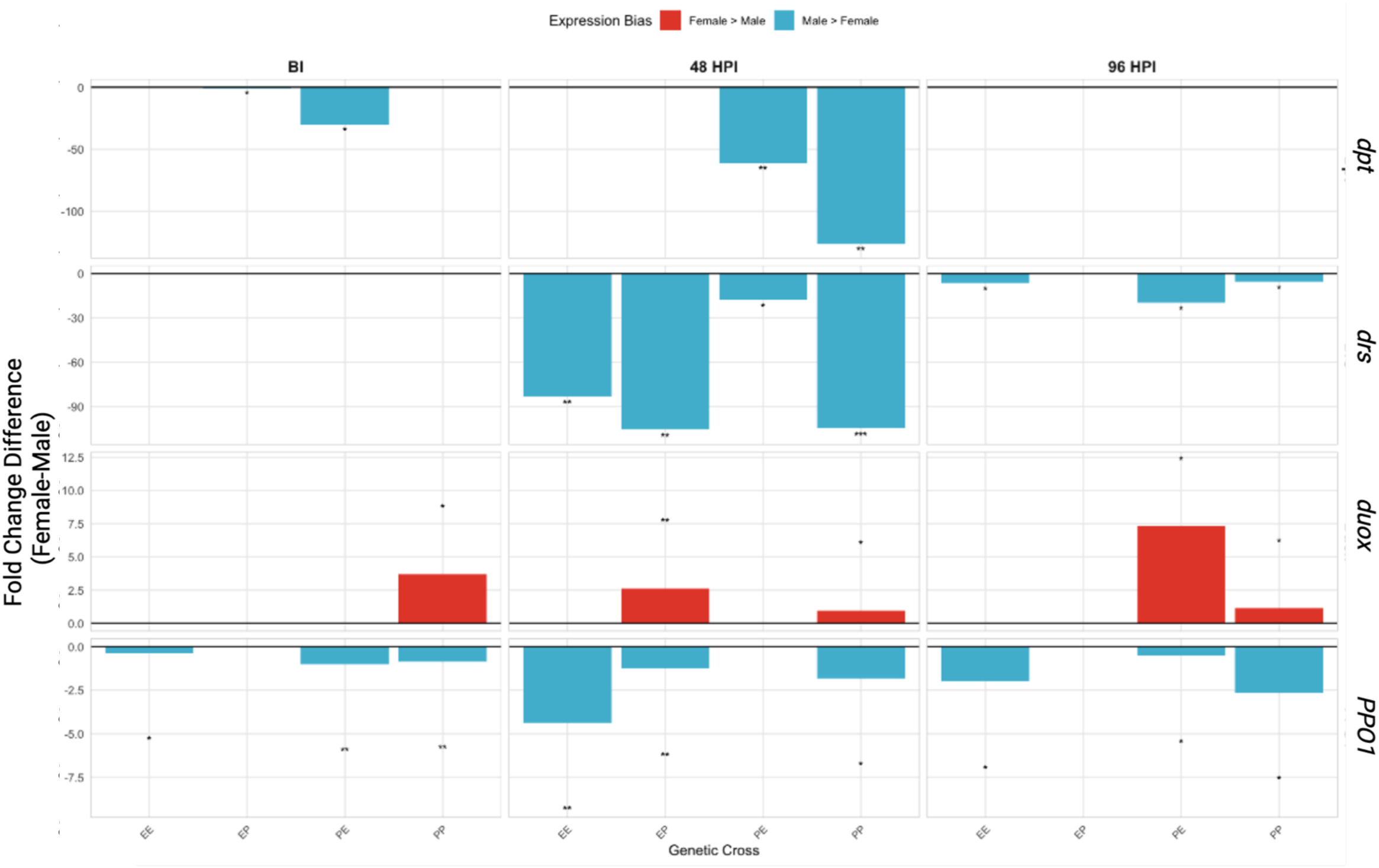
Sexual dimorphism in *Drosophila melanogaster* immune gene expression across genetic backgrounds and infection time points. Bar plots show fold change differences between females and males (Female - Male) for four immune genes (*Duox, Dpt, Drs, PPO1*) measured in four genetic crosses (EE, EP, PE, PP) at three time points: before infection (BI), 48 hours post-infection (48 HPI), and 96 hours post-infection (96 HPI). Positive values (red bars) indicate female-biased expression; negative values (blue bars) indicate male-biased expression. All values are normalized to the control PP Male BI calibrator. Statistical significance is indicated by asterisks: ***p < 0.001, **p < 0.01, *p < 0.05 (Welch’s t-test, n = 3 biological replicates per sex). The data reveal extensive male-biased expression, particularly for antimicrobial peptide genes (*dpt, drs*) during active infection phases.

**Table 3.**
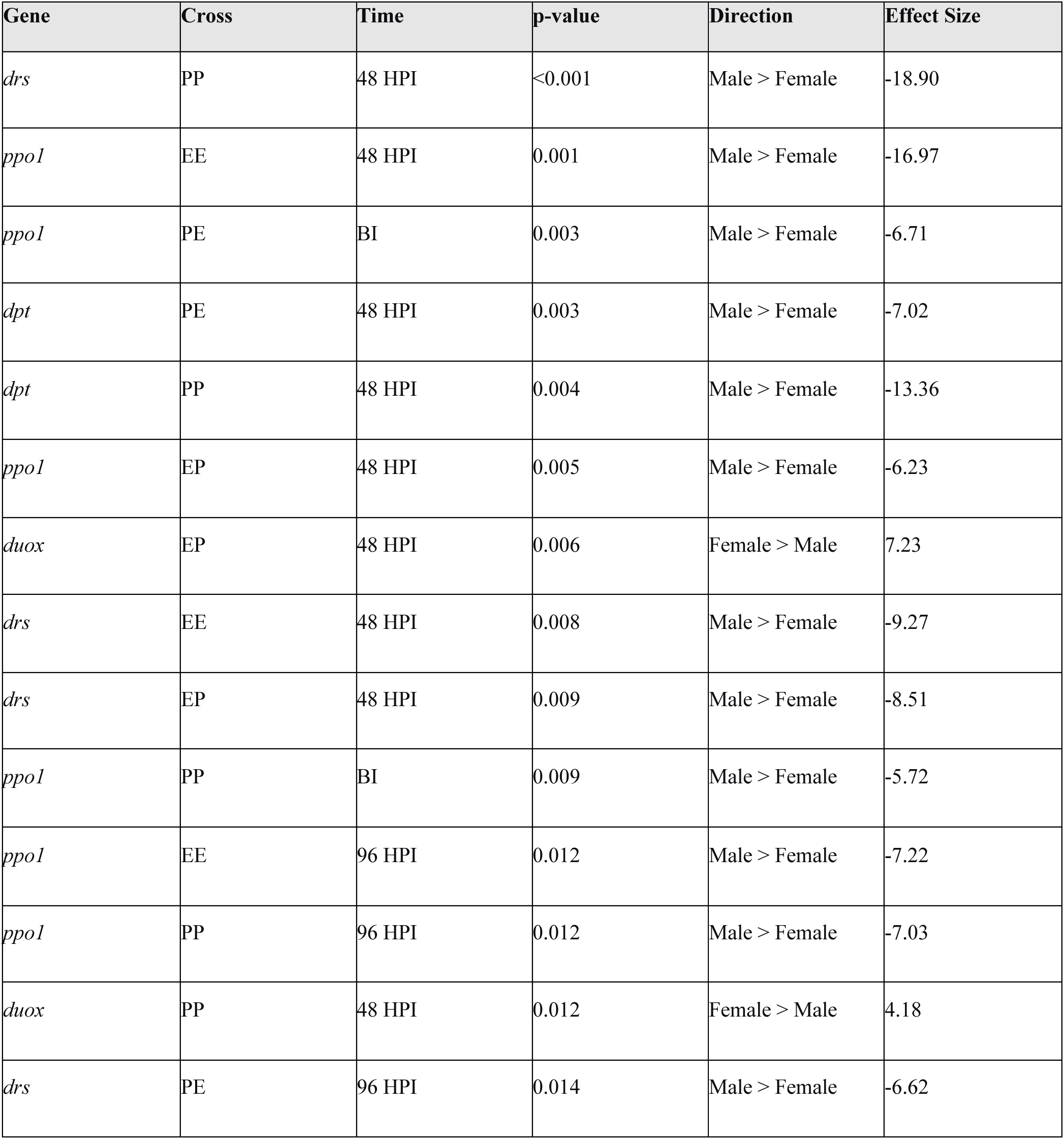
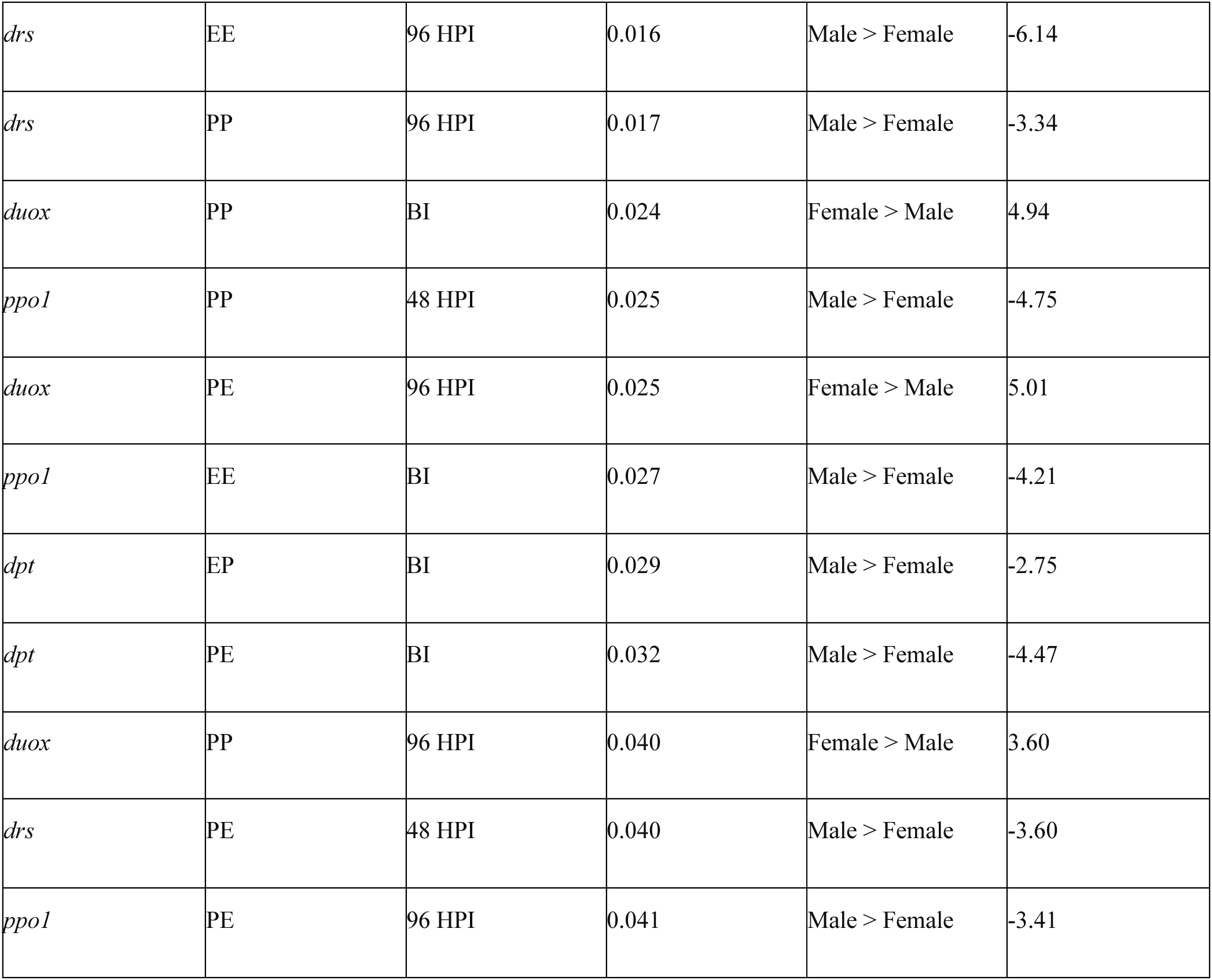
Significant sex differences in immune gene expression across genetic crosses and time points. Results from Welch’s t-tests comparing male and female gene expression (fold change relative to PP Male BI calibrator) for each Gene×Cross×Time combination. Only statistically significant differences (p < 0.05) are shown. Fold change differences represent Female - Male values; positive values indicate female-biased expression, negative values indicate male-biased expression. Effect sizes (Cohen’s d) are classified as: negligible (<0.2), small (0.2-0.5), medium (0.5-0.8), or large (>0.8).

In addition to this sex bias, genetic background effects were sex- and time-dependent. Females showed sensitivity to parental genetic background at earlier stages of infection, whereas males exhibited genetic effects only at the late infection stage (96 HPI), where PE cross showed markedly reduced expression (Figure 5, Table 4).

**Figure 5:**
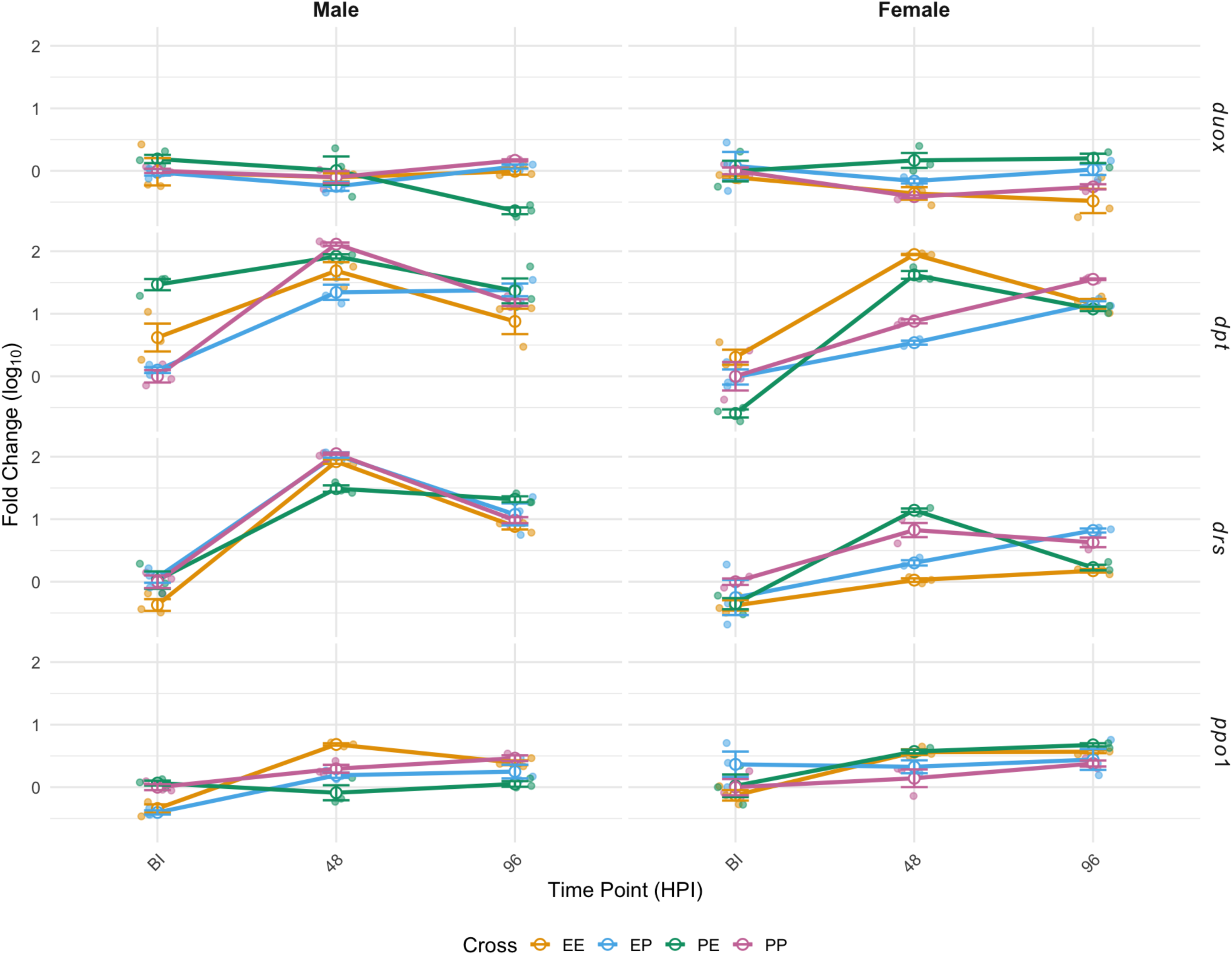
Gene expression dynamics across four F₁ crosses (EE, EP, PE, PP) measured at three time points (BI, 48 HPI, 96 HPI) in females and males. Expression levels (Fold Change) are shown for four immune-related genes (*duox, dpt, drs, ppo1*), with each panel representin1g a unique Sex × Gene combination. Coloured lines represent mean expression trajectories for each cross, with circular points indicating mean values and error bars showing the standard error of the mean (SEM). Light jittered points represent individual biological replicates.

**Table 4.**
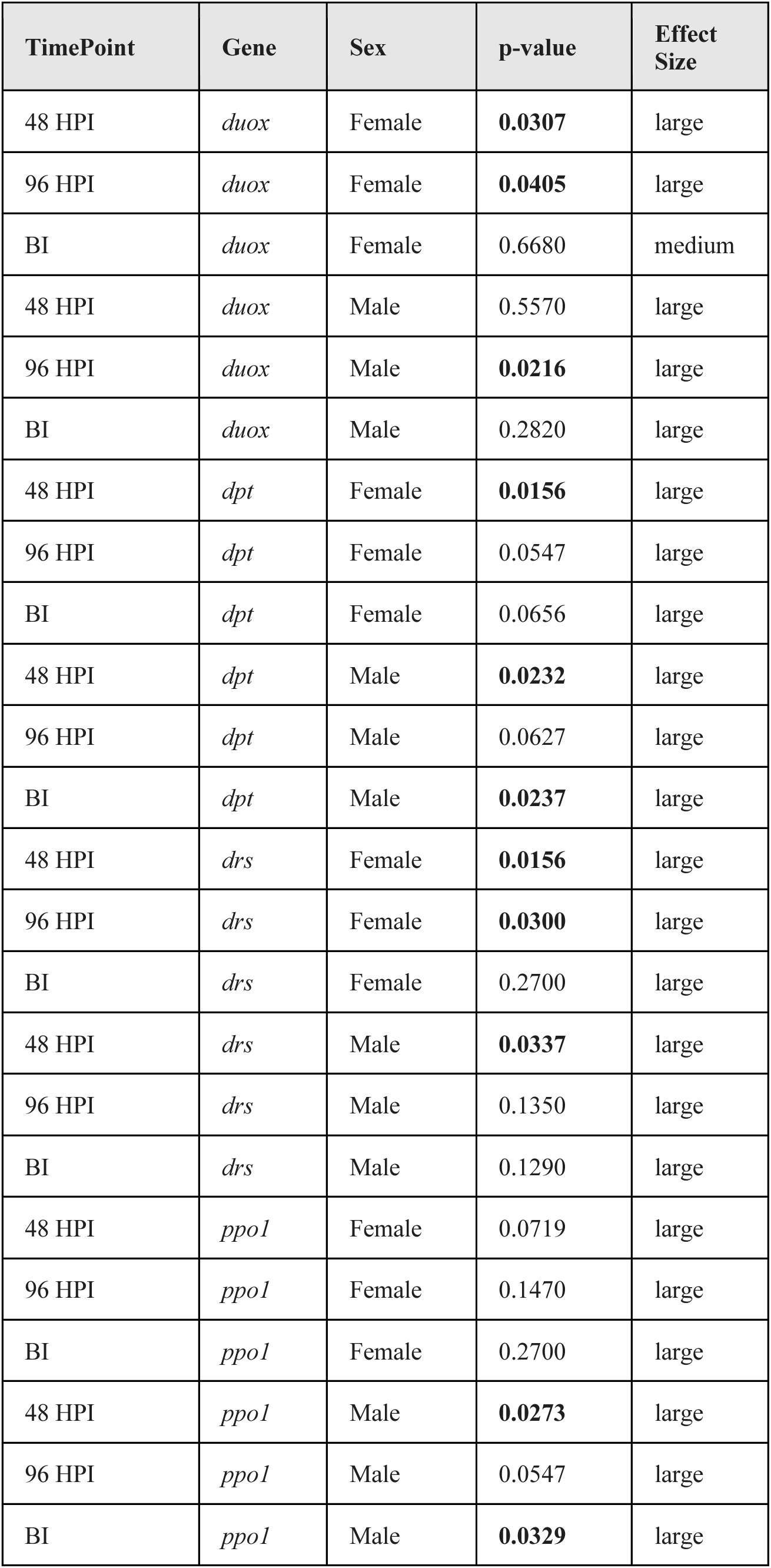
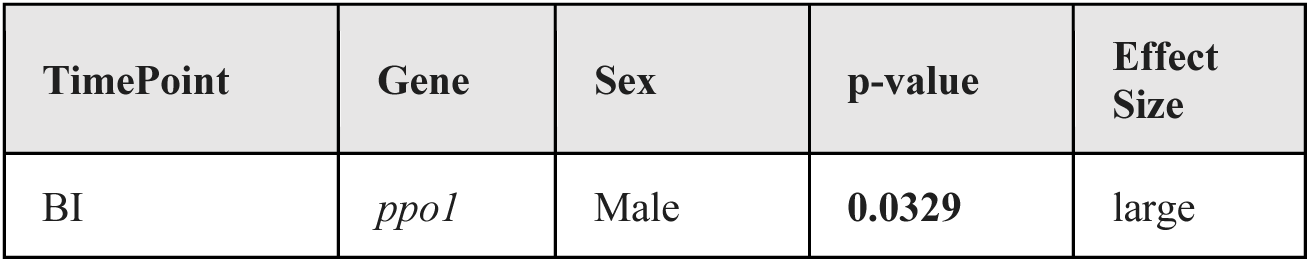
Cross-specific effects on immune gene expression across time points. Results of Kruskal–Wallis tests evaluating whether gene expression differs among F1 crosses (EE, EP, PE, PP) at each time point (BI, 48 HPI, 96 HPI), separately for males and females. For each gene, sex, and time point combination, the table reports the p-value, significance category, and effect size (epsilon-squared; ε²). Significance codes: p < 0.05 = *, p < 0.01 = **, p < 0.001 = ***; ns = not significant. Effect-size categories follow standard ε² interpretations: negligible (<0.01), small (0.01–0.04), medium (0.04–0.16), and large (>0.16).

Temporal dynamics further differed between sexes where females demonstrated a significant downregulation of *duox* during mid-infection, while males showed no clear temporal changes. Together, these results indicate distinct regulatory trajectories of *duox* between sexes, with females mounting earlier, genetically modulated responses and males showing delayed, late-stage genetic effects (Figur 5, Table 5).

**Table 5.**
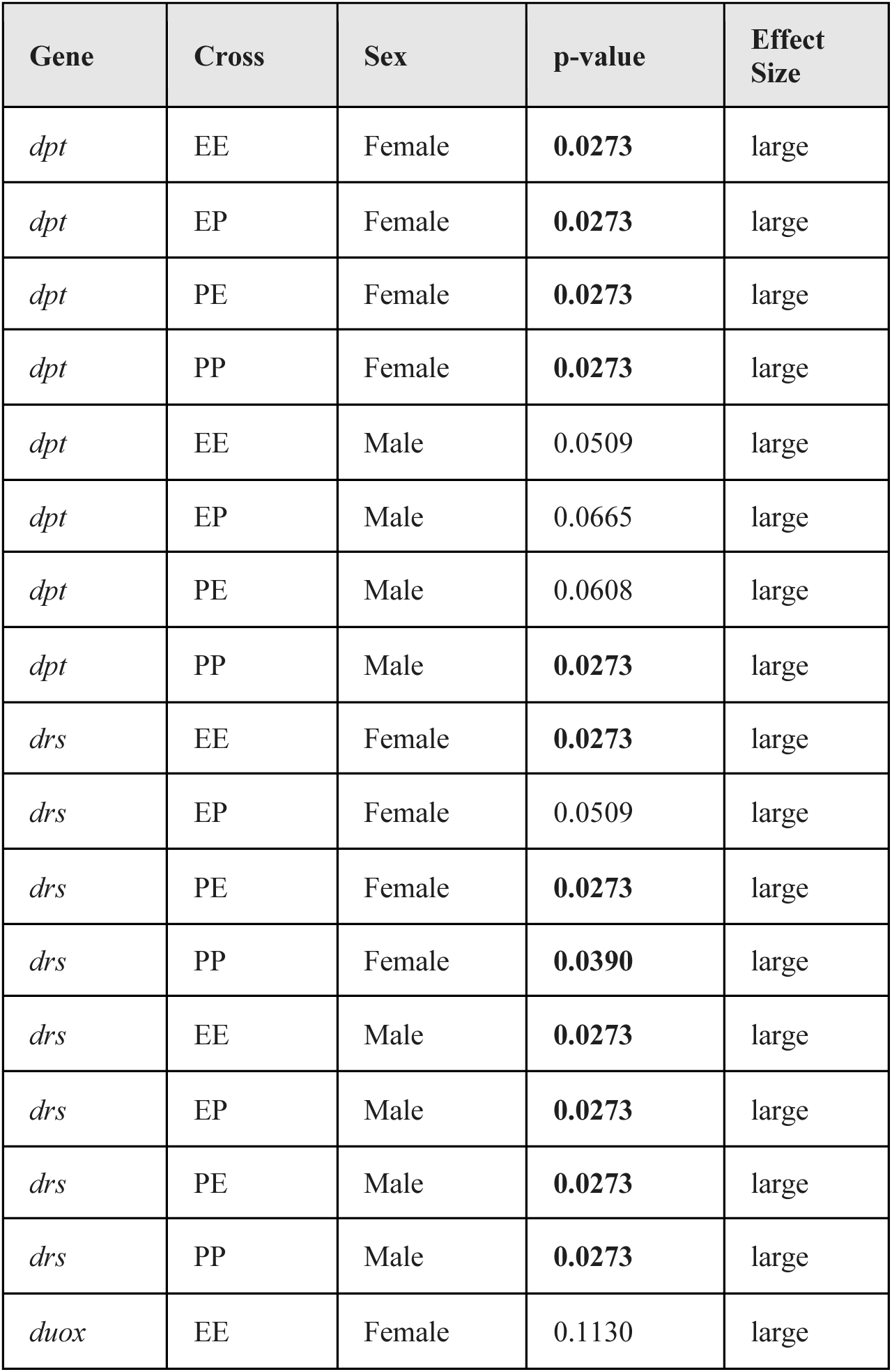
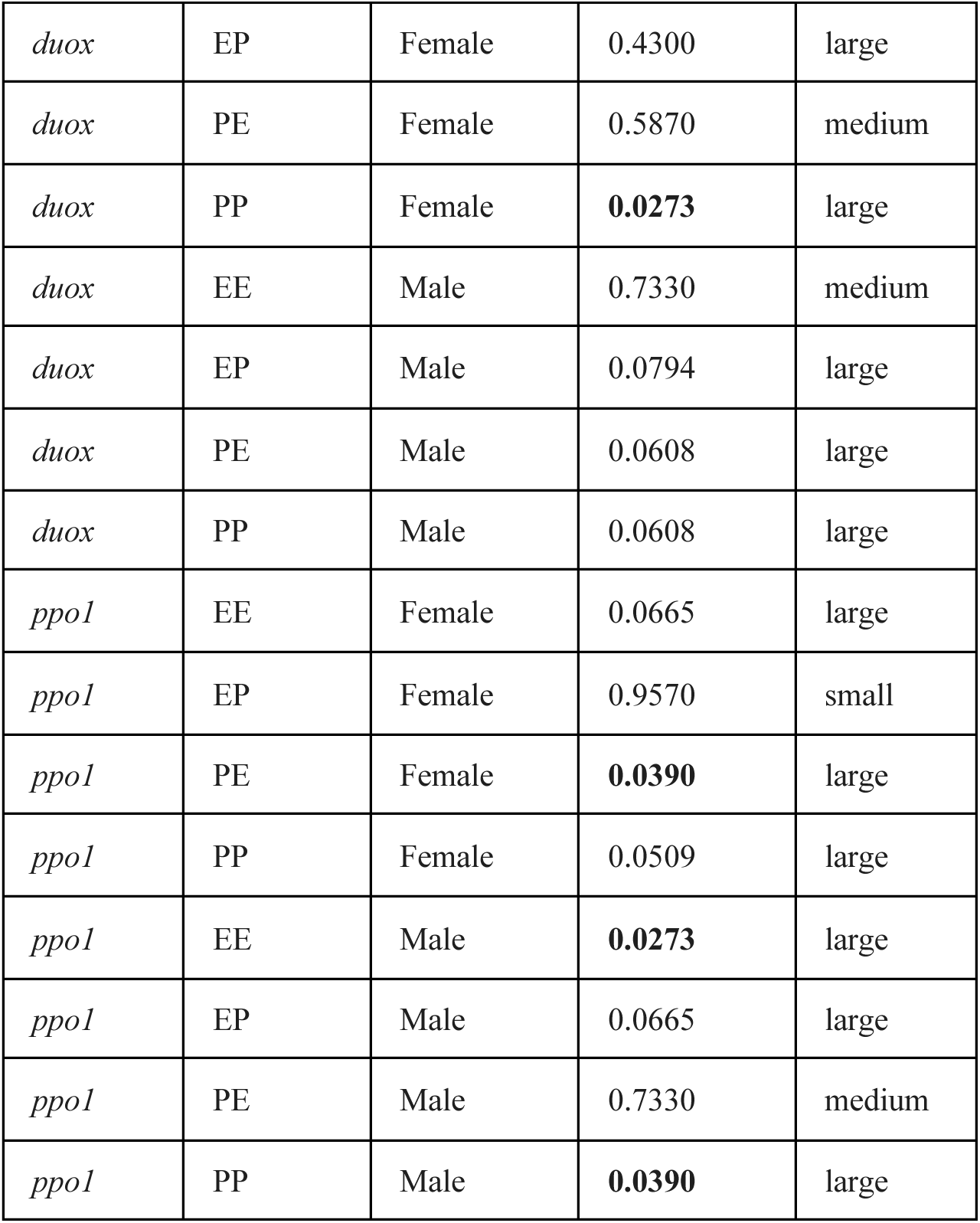
Within-cross temporal changes in immune gene expression across F1 progeny. Kruskal–Wallis tests were performed separately for each immune gene (*duox, dpt, drs, ppo1*), cross (EE, EP, PE, PP), and sex (female, male) to evaluate whether gene expression differed across the three timepoints (BI, 48 HPI, 96 HPI). Each row therefore represents a time-effect test within a specific Sex × Cross × Gene combination. Reported values include the p-value, significance class (p < 0.05 = “*”), and the corresponding effect size (epsilon-squared), categorized as negligible, small, medium, or large.

#### 3.2.2 Diptericin (dpt) shows male-biased expression with widespread temporal induction

*Dpt* exhibited a strong male-biased expression, with males showing substantially higher expression than females across time points (Figure 4, Table 3). Notably, baseline (BI) expression differed significantly among genetic crosses, with PE males displaying elevated basal *dpt* levels compared to other crosses, indicating a strong parent-of-origin effect on constitutive expression prior to infection (Figure 5, Table 4). Following infection, *dpt* was robustly induced in both sexes, peaking at 48 HPI and declining by 96 HPI. While this temporal pattern was conserved across crosses, the magnitude of induction depended on baseline expression, with PE males showing high initial levels and comparatively reduced fold induction (Figure 5, Table 5). In contrast, females exhibited lower baseline expression and more uniform induction across crosses. Together, these results indicate that parental genetic background, particularly in PE crosses, strongly modulates baseline *dpt* expression, which in turn shapes the apparent strength of post-infection induction, especially in males.

#### 3.2.3 Drosomycin (drs) exhibits pronounced male-biased expression with early dominant response

*Drs* exhibited the strongest male-biased expression among all genes examined, with males showing consistently higher expression than females across conditions (Figure 4, Table 3). In addition to this sex bias, *drs* regulation was sexually dimorphic with respect to both genetic background and timing. Males showed significant cross-specific effects exclusively at 48 HPI, indicating early genetic modulation of induction (Figure 5, Table 4). In contrast, females exhibited cross differences at the same time point but between different genetic backgrounds, highlighting sex-specific sensitivity to parental genetic contributions (Figure 5, Table 4). Temporal analyses revealed a robust and conserved early induction of *drs* in males, with significant upregulation at 48 HPI across all genetic crosses (Figure 5, Table 5). Females, however, showed more selective and cross-dependent temporal responses, with some backgrounds responding early and others exhibiting delayed induction at 96 HPI (Figure 5, Table 5).

#### 3.2.4 Prophenoloxidase 1 (PPO1) demonstrates consistent male-biased expression with

*PPO1* expression showed a strong and consistent male bias across most genetic crosses and time points, with males exhibiting significantly higher expression than females under multiple conditions (Figure 4, Table 3). The large effect sizes indicate robust sex-biased regulation of this melanization-related gene. Beyond sex differences, genetic background effects were largely male-specific. Males displayed significant cross-dependent variation at both 48 HPI and 96 HPI, indicating sustained genetic modulation of *PPO1* expression during infection (Figure 5, Table 4). In contrast, females showed no significant cross effects, suggesting fundamentally different regulatory architectures between sexes.

Temporal patterns further emphasized this dimorphism. Males exhibited both early and late induction, depending on genetic background, whereas females showed a single, restricted temporal response, limited to PE crosses at 96 HPI (Figure 5, Table 5). Overall, these results indicate that *PPO1* regulation is predominantly male-driven, with broader temporal flexibility and genetic sensitivity in males compared to tightly constrained regulation in females.

### 3.3. Phenoloxidase activity

We found a significant main effect of Hours post infection (HPI) on log-transformed phenoloxidase (PO) activity. Neither the cross nor sex, had a significant effect (Figure 7, S4; Table 6), and no interactions among factors were significant. The random effect of Block explained negligible variance (SD = 0.0096). These results indicate that PO activity changed over time post-infection but did not differ between treatments or sexes.

**Figure 6:**
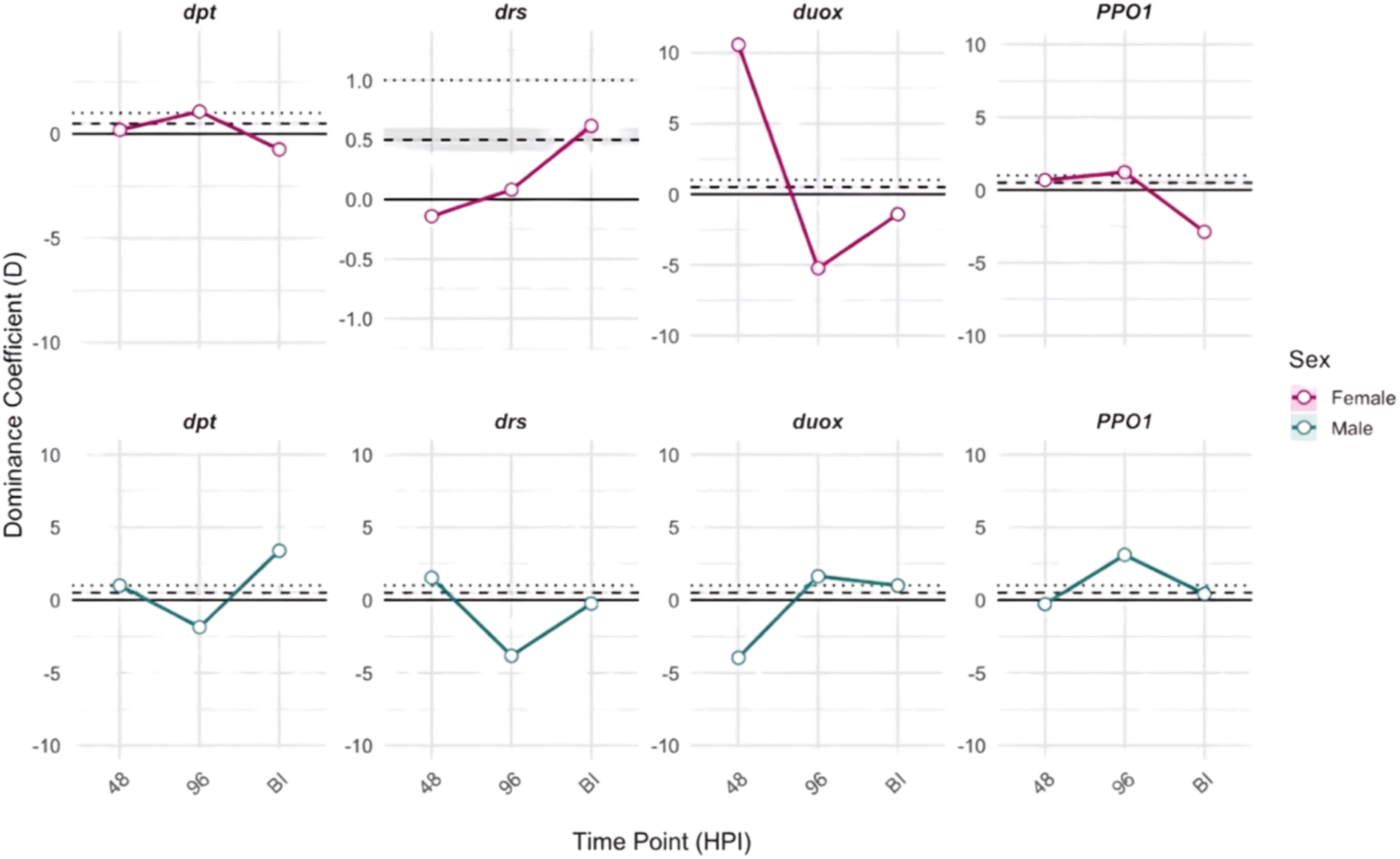
Non-additive inheritance of immune gene expression varies across time and between sexes. Dominance coefficients (D) were calculated for four immune genes (*dpt, drs, duox, PPO1*) at BI, 48 HPI, and 96 HPI. Panels are separated by sex, with each gene shown in a separate column. The solid line marks D = 0 (additive), the dashed line D = 0.5 (partial dominance), and the dotted line D = 1 (complete dominance). Positive D-values indicate dominance toward the higher-expressing homozygote, while negative D-values reflect dominance toward the lower-expressing homozygote. The figure illustrates sex-specific and temporal shifts in dominance relationships following infection. Table corresponding to this figure is provided in supplementary table (Table S2).

**Figure 7.**
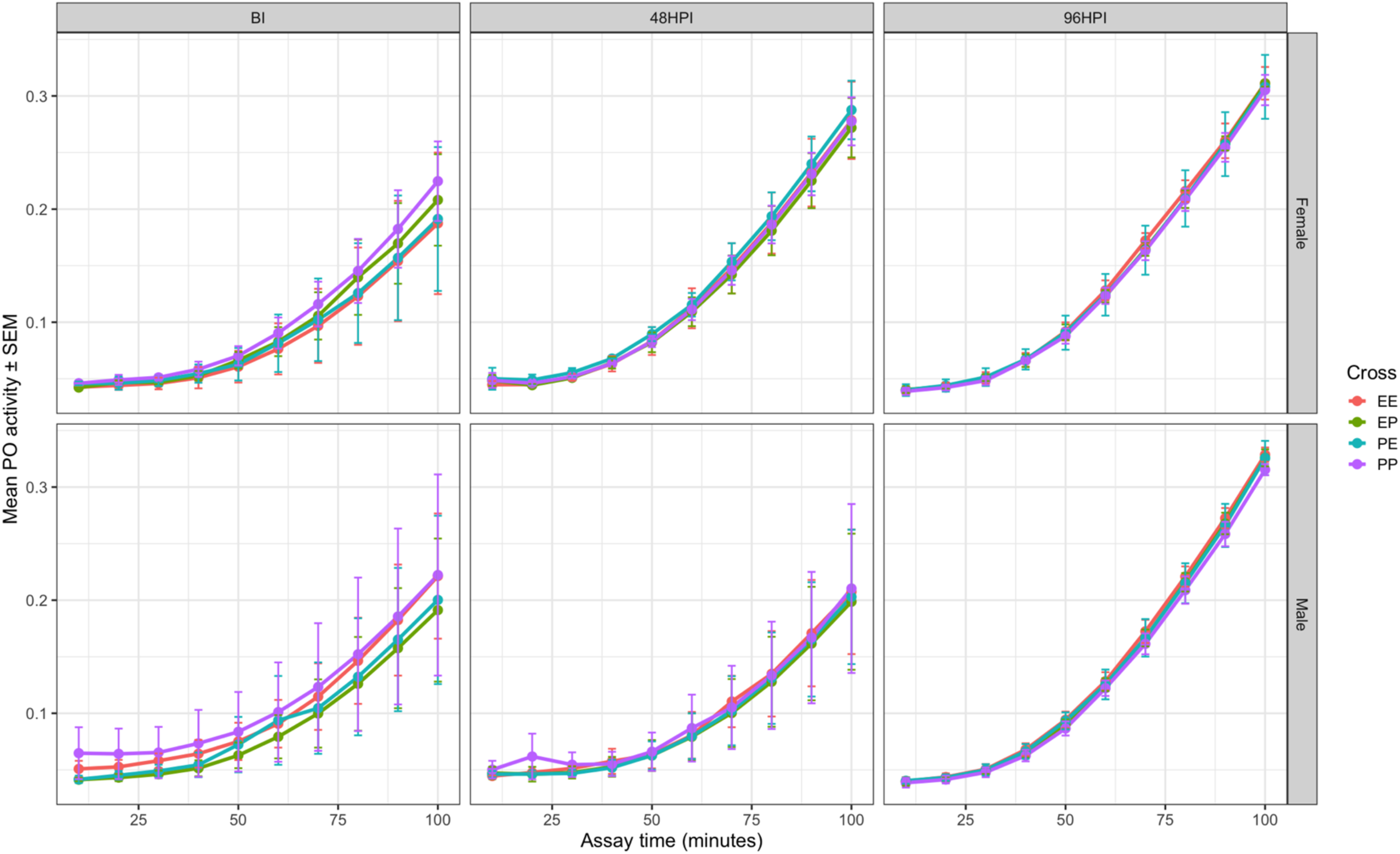
Sex-specific differences in phenoloxidase (PO) activity across infection stages. Mean PO activity (absorbance over assay time) is shown for each Cross (EE, EP, PE, PP), faceted by Sex (rows) and HPI (columns: BI, 48 HPI, 96 HPI). Each panel represents mean absorbance values across biological replicates (Blocks). Block-specific plots are shown in Supplementary Figure S4.

**Table 6.** Type III ANOVA results from the linear mixed-effects model evaluating PO activation rate. The table shows results from a linear mixed-effects model of PO activity slope, where the slope represents the rate of absorbance increase during the PO assay (a proxy for enzyme activation rate). Cross (EE, EP, PE, PP), Sex (Female, Male), and HPI (BI, 48 HPI, 96 HPI) were treated as fixed effects, with Block included as a random intercept. The analysis tests whether PO activation rate differs across genotypes, sexes, infection stages, and their interactions. HPI had a significant main effect, indicating that PO activation rate increases with infection stage, whereas Cross and Sex did not significantly influence slope estimates.

| Effect | Sum Sq | Mean Sq | NumDF | DenDF | F value | Pr(>F) |
| --- | --- | --- | --- | --- | --- | --- |
| Cross | $1.81 \times 10^{-8}$ | $6.00 \times 10^{-9}$ | 3 | 23 | 0.0140 | 0.99768 |
| Sex | $7.98 \times 10^{-7}$ | $7.98 \times 10^{-7}$ | 1 | 23 | 1.8498 | 0.18699 |
| HPI | $1.64 \times 10^{-5}$ | $8.22 \times 10^{-6}$ | 2 | 23 | 19.0646 | **>0.001** |
| Cross:Sex | $6.81 \times 10^{-8}$ | $2.27 \times 10^{-8}$ | 3 | 23 | 0.0526 | 0.98367 |
| Cross:HPI | $1.15 \times 10^{-7}$ | $1.91 \times 10^{-8}$ | 6 | 23 | 0.0443 | 0.999 |
| Sex:HPI | $2.69 \times 10^{-6}$ | $1.35 \times 10^{-6}$ | 2 | 23 | 3.1214 | 0.06319. |
| Cross:Sex:HPI | $1.03 \times 10^{-7}$ | $1.72 \times 10^{-8}$ | 6 | 23 | 0.0399 | 0.99967 |
| Random Effect | Variance | Std. Deviation |  |  |  |  |
| Block | >0.001 | 0.0002 |  |  |  |  |
| Residual | >0.001 | 0.0006 |  |  |  |  |

### 3.4. Fecundity

We found no statistically significant main effects of genetic cross or time period on fecundity (Figure 8, Table 7). However, several patterns approached marginal statistical significance and revealed biologically meaningful trends. PE crosses showed a strong tendency toward higher fecundity compared to EE crosses, laying approximately 65% more eggs during the experimental period. Similarly, the later time period (24-36 hours) showed a marginal increase in fecundity compared to the early period.

**Figure 8.**
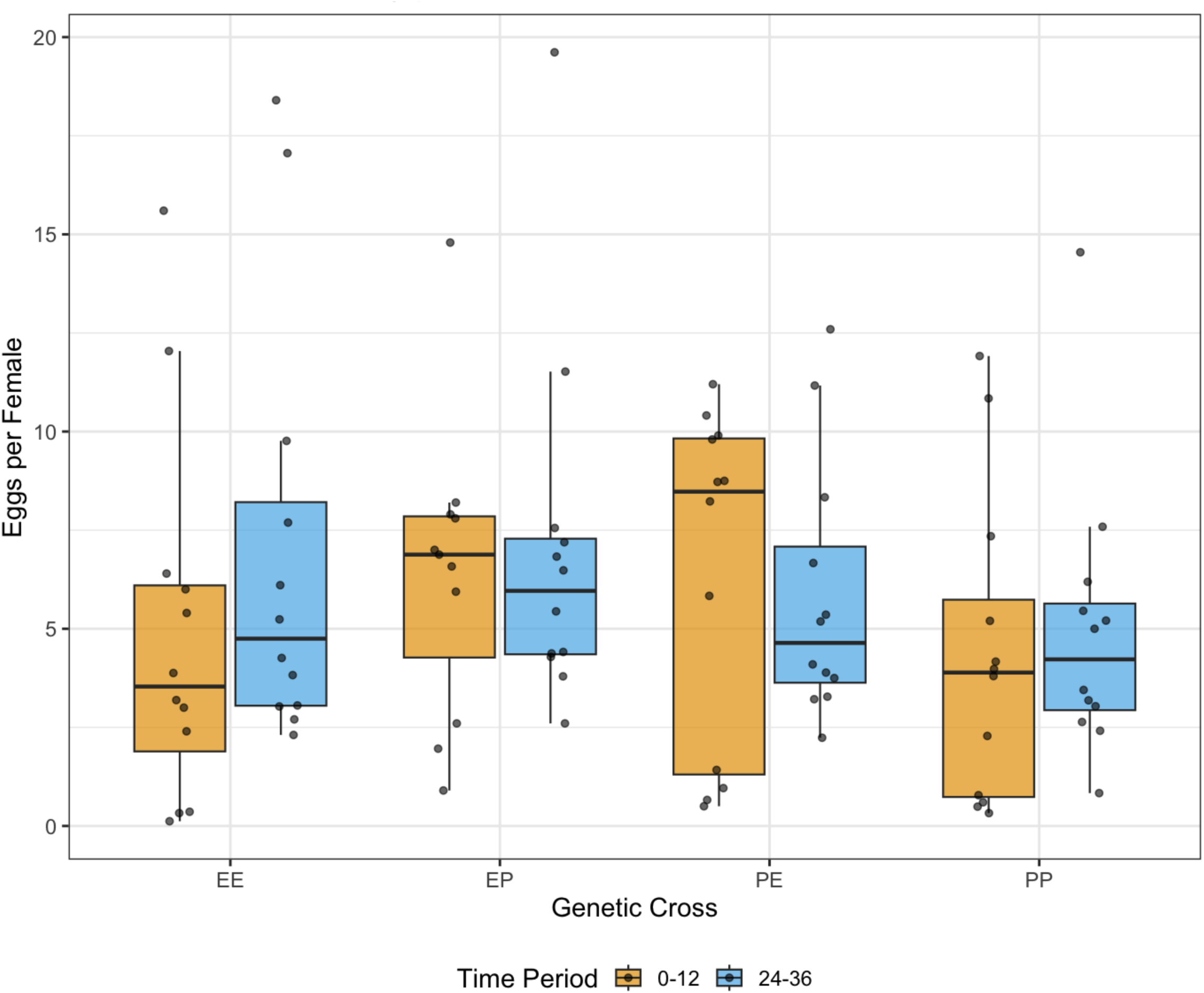
Fecundity distribution by genetic cross and time period post-infection Statistical analysis using negative binomial generalized linear mixed modeling revealed no significant differences between crosses after multiple testing correction (all p > 0.05). Box plots show median (central line), interquartile range (boxes), and data range (whiskers) of eggs laid per surviving female across all experimental blocks. Individual data points are overlaid to show raw data distribution. PE crosses showed the highest median fecundity during the early time period (0-12h), while EP crosses showed the highest median fecundity during the later time period (24-36h).

**Table 7.** Results from negative binomial generalized linear mixed model analyzing fecundity rates. The model included genetic cross (EE, EP, PE, PP), time period (0-12h, 24-36h), and their interaction as fixed effects, with experimental block as a random effect. The response variable was egg counts with an offset for the number of surviving females. While no effects reached statistical significance at α = 0.05, PE crosses showed a tendency toward higher fecundity compared to EE crosses (p = 0.098).

| Fixed Effects | Estimate | Std. Error | z-value | p-value |
| --- | --- | --- | --- | --- |
| Intercept (EE, 0-12h) | 1.395 | 0.274 | 5.093 | <0.001 |
| Cross: EP | 0.393 | 0.304 | 1.294 | 0.196 |
| Cross: PE | 0.504 | 0.305 | 1.654 | 0.098 |
| Cross: PP | 0.000 | 0.299 | 0.001 | 0.999 |
| Time: 24-36h | 0.494 | 0.301 | 1.639 | 0.101 |
| EP × 24-36h | -0.283 | 0.421 | -0.672 | 0.502 |
| PE × 24-36h | -0.677 | 0.422 | -1.602 | 0.109 |
| PP × 24-36h | -0.332 | 0.419 | -0.792 | 0.428 |
| Random Effect | Variance | Std. Deviation |  |  |
| Block | 0.116 | 0.341 |  |  |
| Residual | 1.056 | 1.027 |  |  |

## 4. Discussion

### 4.1. An additive genetic architecture for evolved survivorship, but a complex one for its underlying mechanisms

The objective of this study was to dissect the genetic and physiological basis of evolved post-infection survivorship in *Drosophila melanogaster* following *Enterococcus faecalis* infection. As expected under directional selection, evolved populations (EE) showed the highest survivorship, hybrids (EP and PE) displayed intermediate survival, and controls (PP) showed the lowest. Dominance coefficients for both sexes were close to zero, and hybrid survivorship closely matched additive expectations, indicating that increased survivorship is largely additive in its genetic architecture. The absence of differences between reciprocal hybrids further indicates that this trait is autosomal rather than sex-linked. Together, these results indicate that the increased post infection survivorship in the E population is not due to a few dominant, large-effect alleles, but rather the cumulative effect of many alleles, each making a small, additive contribution to the phenotype—an outcome typical of complex, fitness-related traits (Nelson & Crone, 1999; Hill et al., 2008).

This clear additive pattern observed in survival, however, reflects a more complex situation at the molecular level. When we examined whether differences in gene expression explain the higher survival of EE flies, the answer was not straightforward. Contrary to the expectation that higher post infection survivorship would be explained by uniformly elevated immune gene expression, we found no such simple pattern. Instead, the evolved changes in gene regulation were both sex-specific and time-dependent, indicating that immune regulation differs between males and females and varies over time (Figure 6) (Vincent & Sharp, 2014; Schwenke et al., 2016). For example, while EE flies did not show universally higher AMP expression, they displayed distinct temporal dynamics, such as a delayed but sustained *drs* response in females (Table 5). Our findings reveal specific signatures of selection, such as the significant divergence in *duox* expression between PE and PP males at 96 HPI (Table 5), indicating that evolutionary changes in gene regulation are targeted to specific genetic and temporal contexts. Furthermore, unlike survivorship (which was largely additive), gene expression, showed widespread and often extreme non-additivity. For instance, we observed strong recessivity (e.g., *ppo1* in males at 48 HPI, d = −0.95), strong dominance (e.g., *drs* in females at 48 HPI, d = 0.64), and even overdominance (e.g., *dpt* in males at BI, d = 1.32) (Figure 6, Table S2). These results, taken together, suggest that the superior survivorship of the E population does not result from simply “amping up” the expression of a few key immune genes in a dominant manner. Instead, it most likely results from a more complex, reorganised network of gene regulation that precisely controls the timing, combination, and sex-specific deployment of immune responses and is effective across genetic backgrounds (Lazzaro et al., 2006; Lemaitre & Hoffmann, 2007; Hill et al., 2008).

### 4.2. A Costly Trade-off: Male-Biased Immune Gene Expression Coincides with Reduced Survival

Expanding on this molecular complexity, our study reveals a pronounced sexual dimorphism that is evident at both the organismal and molecular levels, although these differences manifest in opposite directions. Males showed significantly lower survivorship than females (Table 1), yet males consistently exhibited higher expression of effector genes- *drs* and *dpt*, as well as the melanization precursor *PPO1*. The effect sizes were substantial (Table S1) indicating that this male-biased expression pattern is robust across genetic backgrounds.

In contrast, female-biased front-line defense with superior survival was characterized by higher expression of *(duox*), a key enzyme involved in epithelial immunity and reactive oxygen species production. Females achieved better survival outcomes despite showing lower systemic AMP expression. This inverse relationship between transcriptional activity and survival outcome highlights a key sex-specific trade-off in immune strategy. It suggests that the male approach of investing heavily in inducible systemic effectors may be either less efficient or more physiologically costly than the female strategy, which relies on localized epithelial defenses. The pronounced male-biased AMP expression could therefore represent either a compensatory response to other immunological limitations or an energetically expensive investment that contributes to the reduced survivorship observed in males relative to females (Vincent & Sharp, 2014; Gupta et al., 2016; Schwenke et al., 2016). This was further evidenced by sex-specific genetic (cross identity) effects, such as the differential expression of *dpt* between EE and EP females at 48 HPI (Table 5), underscoring that the genetic architecture of immunity is distinct between the sexes. The observed expression patterns are also consistent with the stem cell renewal hypothesis, where lower renewal rates in males could lead to higher persistent bacterial loads, necessitating sustained, high-level AMP expression, whereas more efficient cellular clearance in females could result in different temporal expression profiles (Regan et al., 2016).

Fecundity assays revealed no significant differences in egg-laying rates between crosses or time periods (Figure 8, Table 7), suggesting that evolved post infection survivorship did not come at a detectable cost to short-term reproduction in females. However, subtle or context-dependent fecundity costs might emerge under more stressful or prolonged conditions (McKean & Nunney, 2008) which warrants further investigation.

### 4.3. Time-dependent evolution of gene regulation and the signature of hybrid misregulation

Our temporal analysis reveals that the evolved changes in gene regulation are highly time-dependent. For instance, the significant downregulation of *duox* gene specifically in PE males at 96 HPI is a classic signature of hybrid misregulation. This is not a simple intermediate expression; it is a breakdown of regulation in a specific genetic context (PE), in one sex (males), at a specific time (96 HPI). This pattern suggests that the E and P populations have diverged in their trans-regulatory factors (e.g., transcription factors, signaling pathways) that control *duox*, and that certain combinations of these factors from different parents are incompatible (Landry et al., 2007; McManus et al., 2010). We also identified clear evidence of maternal inheritance, as seen in the significant difference in *drs* expression between EP and PE males at 48 HPI (p = 0.0382; Table 5a), indicating that the parental origin of alleles influences gene expression, likely through cytoplasmic or epigenetic mechanisms (Wolf & Wade, 2016).

### 4.4. The PPO1-PO disconnect: A potential evolutionary constraint on melanization

Despite upregulation of *PPO1* transcripts in some crosses, PO enzymatic activity remained largely unchanged, indicating potent post-transcriptional regulation, resource limitation, or feedback inhibition. Transcript-protein mismatches are well documented and may arise due to translational control or protein stability mechanisms (Takehana et al., 2004). Our results suggest that transcriptional data alone may not adequately capture functional immune capacity and that protein-level assays are essential for assessing immune effector activity. This was particularly evident in males, which showed consistent, significant upregulation of *PPO1* (e.g., in the PP cross at 48 HPI; Table 4a) without a concomitant increase in functional PO activity (Figure 7, S4; Table 6), highlighting a profound sex-specific decoupling of transcription and function.

Additional regulation may occur at the level of *PPO* activation, which requires cleavage by serine proteases and sufficient copper cofactors. Feedback inhibition by serpins may also suppress PO activity (Christensen et al., 2005; Tang et al., 2008). In both honeybees and crustaceans, PO activity varies with nutrition and is often decoupled from *PPO* transcript levels, underscoring the role of metabolic context (Luna-González et al., 2012; Branchiccela et al., 2019). These findings are consistent with the idea that immune activation is energetically costly and tightly regulated (Schmid-Hempel, 2005), particularly in traits like melanization (Nappi & Christensen, 2005).

From an evolutionary standpoint, such tight control over PO activity may reflect the dual role of melanization in defense and self-damage. Melanin synthesis produces cytotoxic compounds and reactive oxygen species that, if unchecked, may harm host tissues (Nappi & Vass, 1993). Therefore, post-transcriptional checkpoints ensure that melanization only occurs when necessary and at appropriate levels. This may also explain why PO activity remained unchanged despite elevated *PPO1* expression; given the potent, cytotoxic potential of the phenoloxidase cascade its activation is tightly constrained in our evolved populations, such that it appears to override evolved changes at the transcriptional level.

These findings highlight the need for an integrative approach such as combining transcriptomics, enzyme assays, and survivorship to accurately assess immune competence (Adamo, 2004). Relying solely on transcriptional data may lead to overestimation of immune potential. Our study demonstrates that a multidimensional perspective is crucial for identifying the specific regulatory bottlenecks, in this case, post-translational control of PPO activation that shape the functional outcome of evolved gene expression changes. This provides a deeper insight into the cost–benefit landscape of immune investment across genetic and sex-specific contexts (Lazzaro & Little, 2009).

## 5. Conclusion

Our study demonstrates that in *Drosophila melanogaster*, the evolved increase in survivorship post infection with *Enterococcus faecalis* is governed by an additive genetic architecture, but the underlying molecular mechanisms are complex, sex-dependent, and temporally dynamic. Compared to females, males have lower survivorship post infection, but show elevated expression of AMP genes. Females rely more on epithelial defence. Thus males and females differ in their post infection survivorship and adopt distinct immune strategies. Our findings reveal a multi-layered adaptive response involving selection line effects, maternal inheritance patterns, and sex-specific regulatory changes. Despite strong molecular divergence, hybrids show additive survivorship, suggesting that regulatory incompatibilities are buffered at the organismal level.

Fecundity analyses further suggest that evolved post infection survivorship does not entail a measurable reproductive cost under laboratory conditions, indicating that the physiological costs of immune investment may manifest primarily through trade-offs other than reduced reproduction. However, future work should test whether such costs appear under resource limitation or multiple infections.

Overall, our findings highlight that adaptation to infection involves not only quantitative changes in immune gene activity but also qualitative shifts in regulatory timing and coordination between sexes. A deeper mechanistic understanding of how these regulatory networks evolve and how they interact with life-history traits like fecundity will provide valuable insights into the balance between immunity, reproduction, and fitness in evolving populations.

## Acknowledgments

We are grateful to Dr. Manas Geeta Arun for his insightful feedback and constructive suggestions, which have significantly strengthened this manuscript. We also acknowledge Devki and Md Kaizer for their expert assistance with the RT-qPCR protocol and for their thoughtful feedback, and the labmates for their support.

## Funding

This work was supported by intramural funding from the Indian Institute of Science Education and Research (IISER) Mohali, India, to N.G.P. T.C. was supported by a Senior Research Fellowship from the Council of Scientific and Industrial Research (CSIR), Government of India. N.K.S. was a postdoctoral fellow funded by IISER Mohali. H.D. was supported by a Senior Research Fellowship from the University Grants Commission (UGC), Government of India. A.D. was supported by IISER Mohali’s institutional Ph.D. fellowship.

## Author Contributions

T.C.: conceptualization, data curation, data analysis, investigation, methodology, validation, writing—original draft, and writing—review and editing.

N.K.S.: methodology, formal analysis, and writing—review and editing.

H.D.: methodology and writing—review and editing.

A.D.: methodology and review.

N.G.P.: conceptualization, formal analysis, methodology, project administration, resources, supervision, and writing—review and editing.

All authors approved the final version of the manuscript and agree to be accountable for all aspects of the work.

## Conflict of Interest

The authors declare no conflict of interest.

## Ethics Statement

This study did not involve human participants or vertebrate animals and did not require approval from an ethics committee.

## Declaration of AI Use

In writing this paper, we did not use any AI-assisted technology.

## Notes

https://dataverse.harvard.edu/previewurl.xhtml?token=754a1c0c-6dfd-4d9d-a4ef-3cf1676eb010

### Competing Interest Statement

The authors have declared no competing interest.

