## Supplementary Figures and Tables for "Non-Additive Transcriptional Regulation Underpins Additive Survival Trait In Populations Of *Drosophila* Selected For Increased Post Infection Survival"

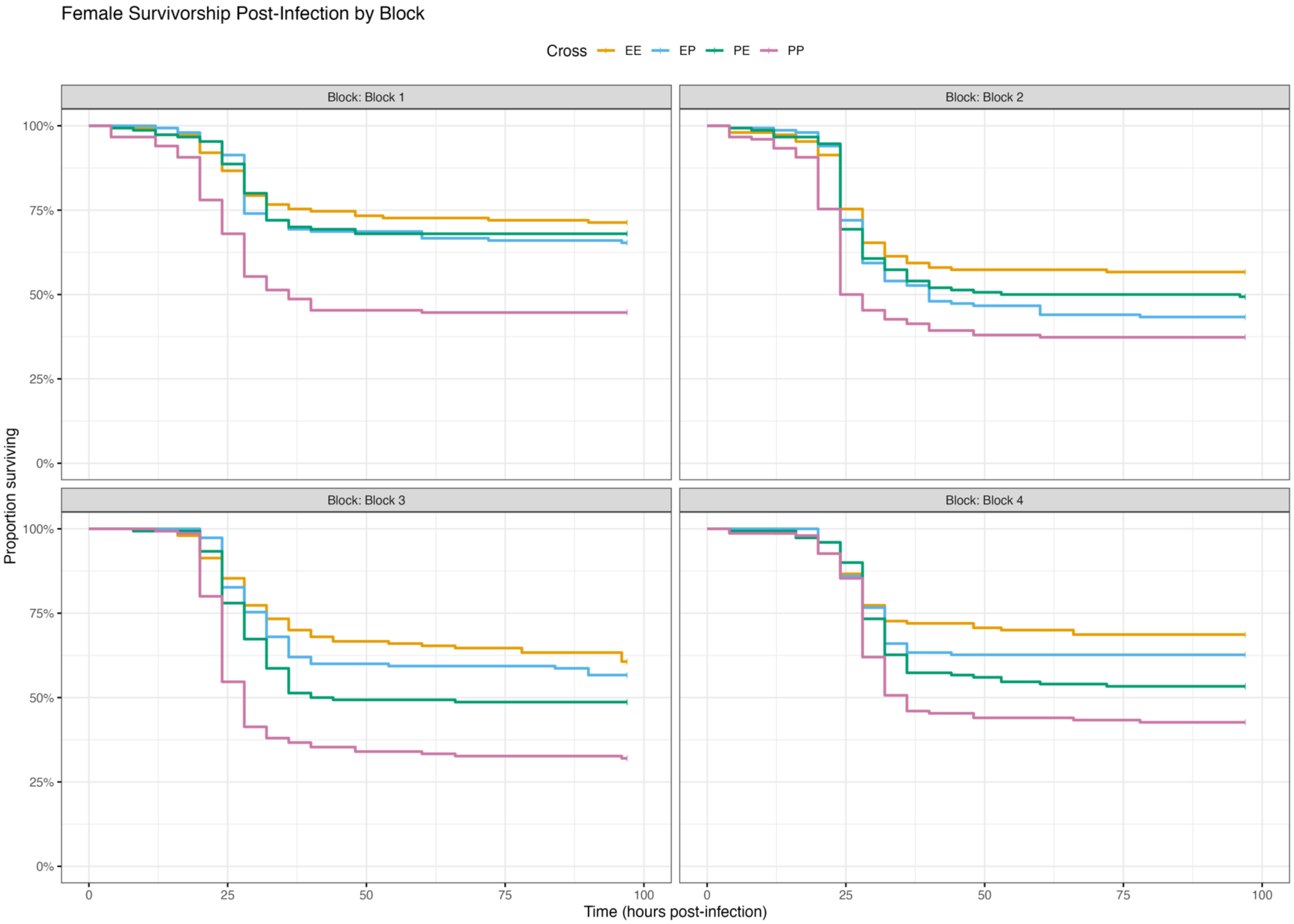


**Supplementary Figure 1**: Survival proportions of Drosophila genetic crosses (EE, PP, EP, PE) across four experimental blocks (1-4) in females. Kaplan-Meier survival curves show temporal survival patterns following infection, with separate panels for each block. Crosses are color-coded as indicated in the legend .

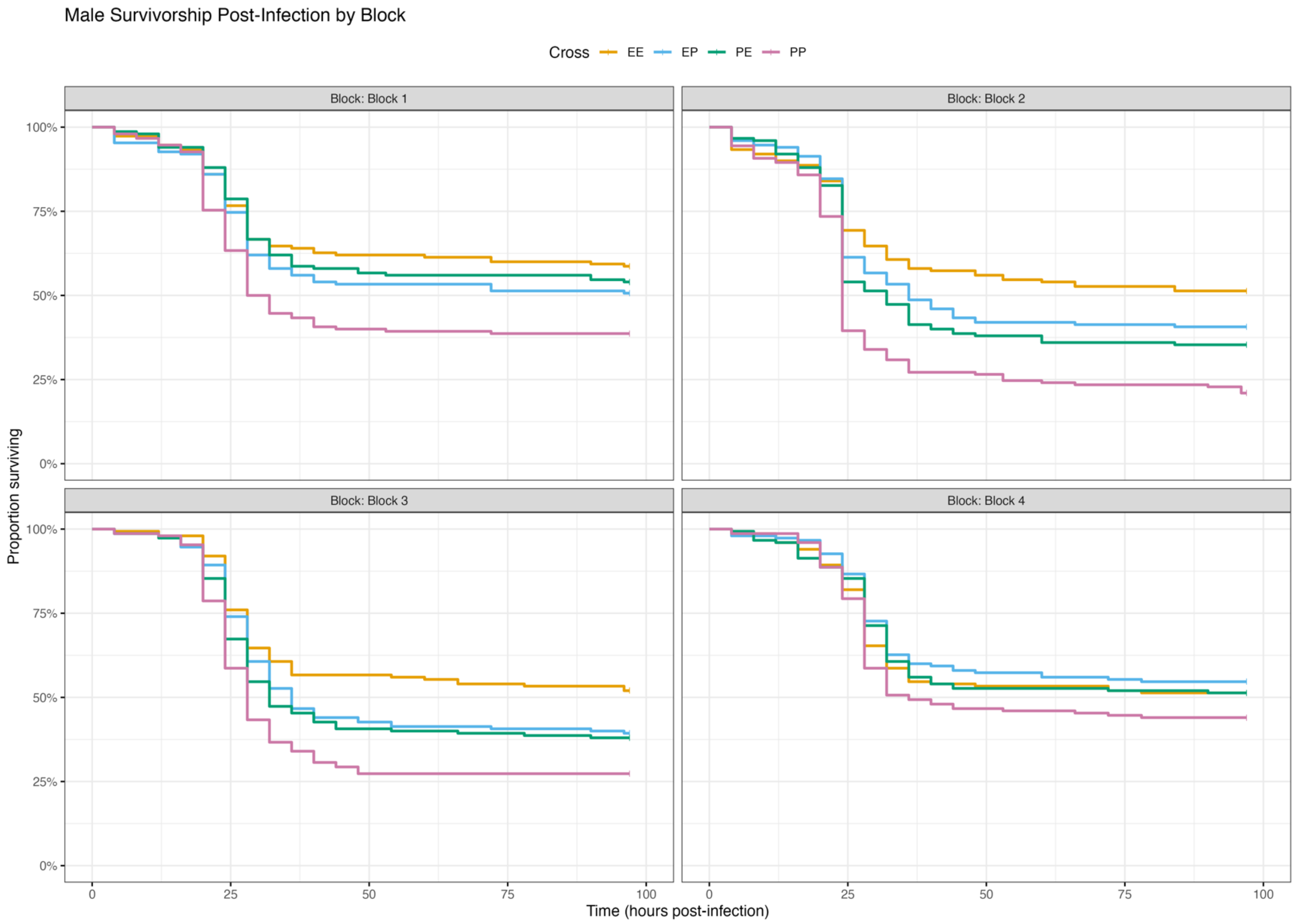


**Supplementary Figure 2**: Survival proportions of Drosophila genetic crosses (EE, PP, EP, PE) across four experimental blocks (1-4) in males. Kaplan-Meier survival curves show temporal survival patterns following infection, with separate panels for each block. Crosses are color-coded as indicated in the legend .

**Supplementary Table 1**: Sex-specific differences in immune gene expression across genetic backgrounds and infection time points. Statistical analysis of fold change differences between females and males for four immune genes (Duox, Dpt, Drs, PPO1) across four genetic crosses (PP, EE, EP, PE) at three time points (BI: before infection; 48 HPI: 48 hours post-infection; 96 HPI: 96 hours post-infection). Values represent fold change differences (Female FC - Male FC) with positive values indicating female-biased expression and negative values indicating male-biased expression. Statistical significance was determined by two-tailed t-test: *$p < 0.05$, **$p < 0.01$, ***$p < 0.001$. Effect sizes were calculated using Cohen's d and categorized as: negligible ($|d| < 0.2$), small ($0.2 \leq |d| < 0.5$), medium ($0.5 \leq |d| < 0.8$), or large ($|d| \geq 0.8$).

| Gene | Cross | Time | Significance | Female FC | Male FC | Difference | Direction | Cohen's d | Effect Size | P-value |
|---|---|---|---|---|---|---|---|---|---|---|
| **Duox** | | | | | | | | | | |
| **Dpt** | PE | 48 HPI | ** | 22.12 | 83.32 | -61.20 | Male > Female | -7.02 | large | 0.00311 |
| **Dpt** | PP | 48 HPI | ** | 3.94 | 130.37 | -126.42 | Male > Female | -13.36 | large | 0.00367 |
| **Dpt** | EP | BI | * | 0.55 | 1.27 | -0.72 | Male > Female | -2.75 | large | 0.02916 |
| **Dpt** | PE | BI | * | 0.13 | 30.41 | -30.28 | Male > Female | -4.47 | large | 0.03175 |
| **Drs** | EE | 48 HPI | ** | 1.07 | 84.02 | -82.96 | Male > Female | -9.27 | large | 0.00766 |
| **Dpt** | | | | | | | | | | |
| **Drs** | EP | 48 HPI | ** | 2.02 | 107.32 | -105.30 | Male > Female | -8.51 | large | 0.00905 |
| **Drs** | PE | 48 HPI | * | 13.98 | 31.36 | -17.38 | Male > Female | -3.60 | large | 0.03989 |
| **Drs** | PP | 48 HPI | *** | 7.14 | 111.90 | -104.76 | Male > Female | -18.90 | large | < 0.001 |
| **Drs** | EE | 96 HPI | * | 1.50 | 7.76 | -6.26 | Male > Female | -6.14 | large | 0.01599 |
| **Drs** | | | | | | | | | | |
| **Drs** | PE | 96 HPI | * | 1.70 | 21.10 | -19.39 | Male > Female | -6.62 | large | 0.01436 |
| **Drs** | PP | 96 HPI | * | 4.40 | 9.77 | -5.37 | Male > Female | -3.34 | large | 0.01677 |
| **Duox** | EP | 48 HPI | ** | 3.20 | 0.58 | 2.62 | Female > Male | 7.23 | large | 0.00599 |
| **Duox** | PP | 48 HPI | * | 1.79 | 0.83 | 0.96 | Female > Male | 4.18 | large | 0.01237 |
| **Duox** | PE | 96 HPI | * | 7.52 | 0.23 | 7.29 | Female > Male | 5.01 | large | 0.02549 |
| **Duox** | PP | 96 HPI | * | 2.59 | 1.47 | 1.11 | Female > Male | 3.60 | large | 0.03967 |
| **Duox** | PP | BI | * | 4.69 | 1.01 | 3.69 | Female > Male | 4.94 | large | 0.02423 |
| **PPO1** | | | | | | | | | | |
| **PPO1** | EE | 48 HPI | ** | 0.47 | 4.83 | -4.36 | Male > Female | -16.97 | large | 0.00128 |
| **PPO1** | EP | 48 HPI | ** | 0.29 | 1.56 | -1.27 | Male > Female | -6.23 | large | 0.00512 |
| **PPO1** | PP | 48 HPI | * | 0.20 | 2.01 | -1.82 | Male > Female | -4.75 | large | 0.02493 |
| **PPO1** | EE | 96 HPI | * | 0.47 | 2.48 | -2.01 | Male > Female | -7.22 | large | 0.01160 |
| **PPO1** | PE | 96 HPI | * | 0.61 | 1.14 | -0.53 | Male > Female | -3.41 | large | 0.04075 |
| **PPO1** | PP | 96 HPI | * | 0.31 | 2.94 | -2.63 | Male > Female | -7.03 | large | 0.01220 |
| **PPO1** | EE | BI | * | 0.10 | 0.47 | -0.37 | Male > Female | -4.21 | large | 0.02651 |
| **PPO1** | PE | BI | ** | 0.16 | 1.17 | -1.01 | Male > Female | -6.71 | large | 0.00256 |
| **PPO1** | PP | BI | ** | 0.14 | 1.01 | -0.87 | Male > Female | -5.72 | large | 0.00938 |

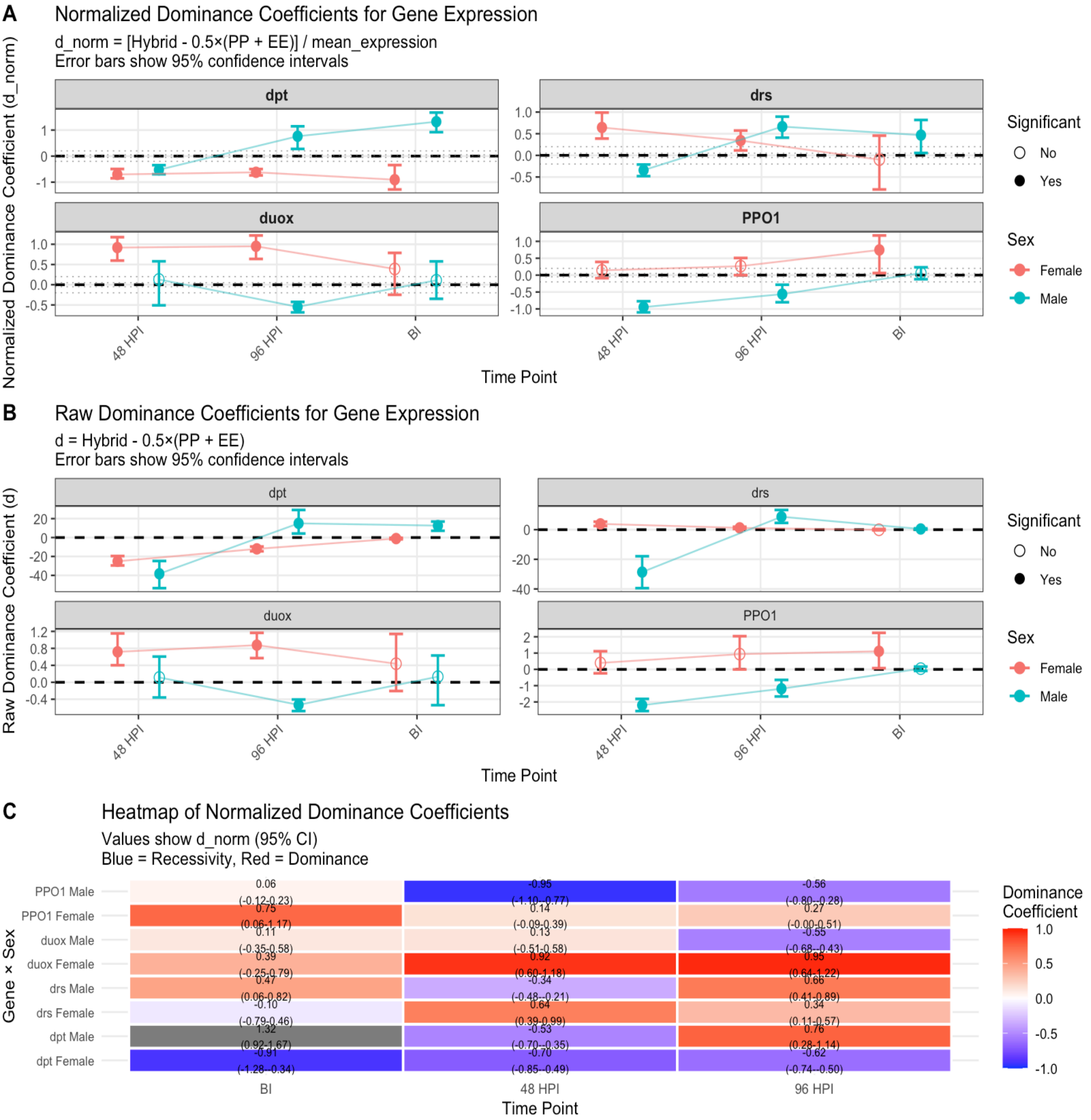
A
Normalized Dominance Coefficients for Gene Expression
d_norm = [Hybrid - 0.5×(PP + EE)] / mean_expression
Error bars show 95% confidence intervals
Normalized Dominance Coefficient (d_norm)
dpt
drs
duox
PPO1
48 HPI
96 HPI
BI
Time Point
Significant
No
Yes
Sex
Female
Male
B
Raw Dominance Coefficients for Gene Expression
d = Hybrid - 0.5×(PP + EE)
Error bars show 95% confidence intervals
Raw Dominance Coefficient (d)
dpt
drs
duox
PPO1
48 HPI
96 HPI
BI
Time Point
Significant
No
Yes
Sex
Female
Male
C
Heatmap of Normalized Dominance Coefficients
Values show d_norm (95% CI)
Blue = Recessivity, Red = Dominance
Gene × Sex
PPO1 Male
PPO1 Female
duox Male
duox Female
drs Male
drs Female
dpt Male
dpt Female
0.06 (-0.12-0.23)
0.75 (0.06-1.17)
0.11 (-0.35-0.58)
0.39 (-0.25-0.79)
0.47 (0.06-0.82)
-0.10 (-0.79-0.46)
1.32 (0.92-1.67)
-0.91 (-1.28--0.34)
-0.95 (-1.10--0.77)
0.14 (-0.09-0.39)
0.13 (-0.51-0.58)
0.92 (0.60-1.18)
-0.34 (-0.48--0.21)
0.64 (0.39-0.99)
-0.53 (-0.70--0.35)
-0.70 (-0.85--0.49)
-0.56 (-0.80--0.28)
0.27 (-0.00-0.51)
-0.55 (-0.68--0.43)
0.95 (0.64-1.22)
0.66 (0.41-0.89)
0.34 (0.11-0.57)
0.76 (0.28-1.14)
-0.62 (-0.74--0.50)
BI
48 HPI
96 HPI
Time Point
Dominance Coefficient
1.0
0.5
0.0
-0.5
-1.0

**Supplementary Figure 3**: Gene expression dominance in hybrids relative to evolved (EE) and control (PP) populations. **(A) Normalized Coefficients (d_norm):** $[Hybrid - 0.5 \times (PP + EE)]/mean\_expression$. Values near 0 are additive; >0 indicates E allele dominance; <0 indicates E allele recessivity. **(B) Raw Coefficients (d):** $Hybrid - 0.5 \times (PP + EE)$, showing absolute expression difference. In A & B, points are jittered; error bars are 95% CI. Filled points are statistically significant. **(C) Heatmap of d_norm:** Visual summary of coefficients from panel A. Red indicates dominance; blue indicates recessivity. Values and 95% CIs are shown.

**Supplementary Table S2: Summary of dominance coefficients for immune gene expression.**

The table presents raw (d) and normalized (d_norm) dominance coefficients for four immune genes across sexes and time points (BI, 48 HPI, 96 HPI)

| Gene | Sex | Time | d_point | d_ci_lower | d_ci_upper | d | mean_expression | d_norm | interpretation | significant |
|---|---|---|---|---|---|---|---|---|---|---|
| **duox** | Male | BI | 0.133 | -0.539 | 0.631 | 0.133 | 1.205 | 0.110 | Moderate Dominance | No |
| **duox** | Male | 48 HPI | 0.114 | -0.358 | 0.607 | 0.114 | 0.874 | 0.130 | Moderate Dominance | No |
| **duox** | Male | 96 HPI | -0.532 | -0.679 | -0.406 | -0.532 | 0.972 | -0.548 | Strong Recessivity | Yes |
| **dpt** | Male | BI | 12.606 | 7.156 | 16.897 | 12.606 | 9.539 | 1.322 | Strong Dominance | Yes |
| **dpt** | Male | 48 HPI | -38.291 | -53.543 | -24.757 | -38.291 | 72.755 | -0.526 | Strong Recessivity | Yes |
| **dpt** | Male | 96 HPI | 15.013 | 4.267 | 29.186 | 15.013 | 19.695 | 0.762 | Strong Dominance | Yes |
| **drs** | Male | BI | 0.456 | 0.050 | 0.882 | 0.456 | 0.977 | 0.467 | Strong Dominance | Yes |
| **drs** | Male | 48 HPI | -28.622 | -39.492 | -17.940 | -28.622 | 83.651 | -0.342 | Strong Recessivity | Yes |
| **drs** | Male | 96 HPI | 8.717 | 4.590 | 13.314 | 8.717 | 13.127 | 0.664 | Strong Dominance | Yes |
| **PPO1** | Male | BI | 0.045 | -0.093 | 0.172 | 0.045 | 0.761 | 0.059 | Moderate Dominance | No |
| **PPO1** | Male | 48 HPI | -2.199 | -2.561 | -1.816 | -2.199 | 2.322 | -0.947 | Strong Recessivity | Yes |
| **PPO1** | Male | 96 HPI | -1.189 | -1.666 | -0.647 | -1.189 | 2.114 | -0.563 | Strong Recessivity | Yes |
| **duox** | Female | BI | 0.438 | -0.205 | 1.139 | 0.438 | 1.121 | 0.390 | Strong Dominance | No |
| **duox** | Female | 48 HPI | 0.718 | 0.402 | 1.152 | 0.718 | 0.782 | 0.918 | Strong Dominance | Yes |
| **duox** | Female | 96 HPI | 0.876 | 0.570 | 1.166 | 0.876 | 0.921 | 0.951 | Strong Dominance | Yes |
| **dpt** | Female | BI | -1.089 | -1.879 | -0.318 | -1.089 | 1.203 | -0.906 | Strong Recessivity | Yes |
| **dpt** | Female | 48 HPI | -25.097 | -29.501 | -19.537 | -25.097 | 35.653 | -0.704 | Strong Recessivity | Yes |
| **dpt** | Female | 96 HPI | -11.982 | -14.516 | -9.504 | -11.982 | 19.285 | -0.621 | Strong Recessivity | Yes |
| **drs** | Female | BI | -0.068 | -0.425 | 0.431 | -0.068 | 0.689 | -0.099 | Moderate Recessivity | No |
| **drs** | Female | 48 HPI | 3.897 | 2.486 | 5.371 | s | 6.053 | 0.644 | Strong Dominance | Yes |
| **drs** | Female | 96 HPI | 1.216 | 0.437 | 1.965 | 1.216 | 3.561 | 0.341 | Strong Dominance | Yes |
| **PPO1** | Female | BI | 1.110 | 0.075 | 2.244 | 1.110 | 1.489 | 0.746 | Strong Dominance | Yes |
| **PPO1** | Female | 48 HPI | 0.396 | -0.241 | 1.113 | 0.396 | 2.795 | 0.142 | Moderate Dominance | No |
| **PPO1** | Female | 96 HPI | 0.936 | -0.003 | 2.037 | 0.936 | 3.527 | 0.265 | Strong Dominance | No |

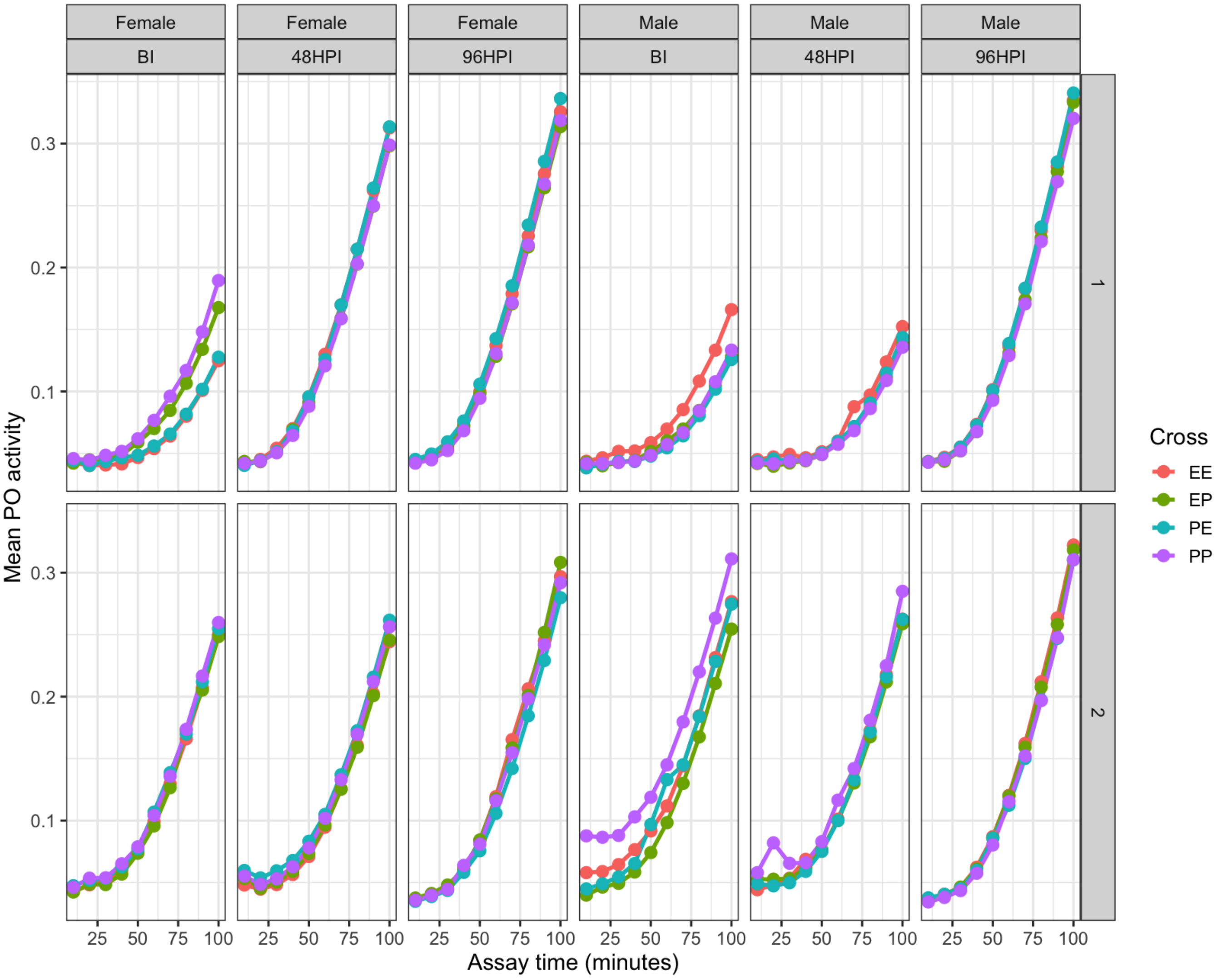


**Supplementary Figure S4:** Phenoloxidase (PO) activity dynamics across genetic crosses, sexes, infection time points, and experimental blocks.

Mean PO activity is shown as a function of assay time (minutes) for the four genetic crosses (EE, EP, PE, PP), plotted separately for females and males and for each infection time point (BI, 48 HPI, 96 HPI). Panels are faceted by sex and HPI, with experimental blocks displayed as separate rows (Block 1 on top, Block 2 on bottom). Points represent mean absorbance values at each assay time within a block, and lines depict the temporal progression of PO activity. This representation allows direct visual comparison of PO kinetics across blocks while preserving sex- and time-point–specific patterns.

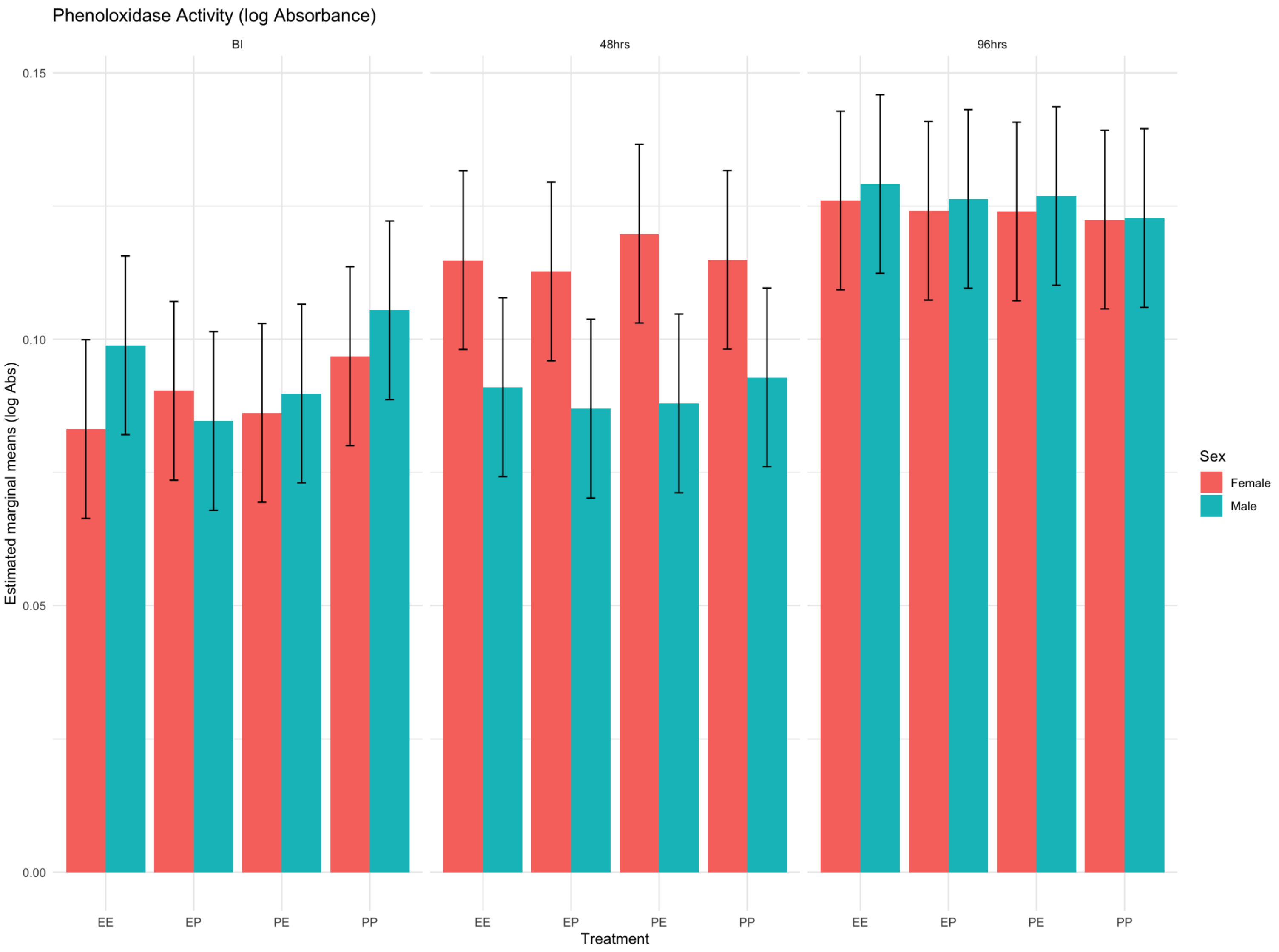


**Supplementary Figure S5**: Phenoloxidase activity (log absorbance) across crosses, sexes, and time points.

Estimated marginal means (± SE) of phenoloxidase (PO) activity are shown for four genetic crosses (EE, EP, PE, PP) at three time points: before infection (BI), 48 hours post-infection, and 96 hours post-infection. Bars represent females (red) and males (teal). Across all crosses, PO activity increases with time since infection, with the highest activity observed at 96 hours.
